# Functional and Structural Characterization of F_1_-ATPase with common ancestral core domains in stator ring

**DOI:** 10.1101/2025.04.30.651372

**Authors:** Aya K. Suzuki, Ryutaro Furukawa, Meghna Sobti, Simon H. J. Brown, Alastair G. Stewart, Satoshi Akanuma, Hiroshi Ueno, Hiroyuki Noji

**Affiliations:** Department of Life Sciences, Graduate School of Arts and Sciences, The University of Tokyo, Tokyo, Japan; Department of Applied Chemistry, Graduate School of Engineering, The University of Tokyo, Tokyo, Japan; Molecular, Structural and Computational Biology Division, The Victor Chang Cardiac Research Institute, NSW, Australia; St Vincent’s Clinical School, Faculty of Medicine, UNSW Sydney, NSW, Australia; School of Science, Molecular Horizons, and the Australian Research Council Centre for Cryo-electron Microscopy of Membrane Proteins, University of Wollongong, Wollongong, NSW, Australia; Faculty of Human Sciences, Waseda University, Saitama, Japan; Research Institute of Planetary Health (RIPH), The University of Tokyo, Tokyo, Japan

## Abstract

Extant F_1_-ATPases exhibit diverse rotational stepping behaviors—3-, 6-, or 9-step cycles—yet the evolutionary origin of these patterns remains unclear. Here, we used ancestral sequence reconstruction to infer the catalytic β and non-catalytic α subunits of a putative ancestral F_1_-ATPase. We then fused their functionally critical domains into the thermostable F_1_ from *Bacillus* PS3, yielding a stable chimeric enzyme. Cryo-EM revealed two distinct conformational states—binding and catalytic dwell states—separated by a ∼34° rotation of the γ subunit, suggesting a fundamental six-step mechanism akin to that of extant 6-stepping F_1_-ATPases. Single-molecule rotation assays with ATP and the slowly hydrolyzed ATP analog ATPγS demonstrated that the chimeric motor is intrinsically a 6-stepper, pausing at binding and catalytic dwell positions separated by 32.1°, although the binding dwell is significantly prolonged by an unknown mechanism. These findings indicate that F_1_-ATPase was originally a 6-stepper and diversified into 3-, 6- and 9-step forms in evolutional adaptation. Based on these results, we discuss plausible features of the entire F_o_F_1_ complex, along with potential physiological contexts in last universal common ancestor and related lineages.

## Introduction

F-type ATP synthase catalyzes the terminal reaction of oxidative phosphorylation—namely, the synthesis of ATP from ADP and inorganic phosphate (P_i_) driven by proton translocation down a proton motive force (*pmf*) across biomembranes. This enzyme is among the most ubiquitous in nature, found in the plasma membranes of prokaryotic cells, the thylakoid membranes of chloroplasts, and the inner membranes of mitochondria. Recent comparative genomic analyses have revealed that cells of the last universal common ancestor (LUCA) already possessed F-type or relevant type ATP synthase[1][2]. These findings provide significant insights into the metabolic capabilities and ecological contexts of LUCA and its descendants, including the last bacterial common ancestor (LBCA) and the last archaeal common ancestor (LACA)[1][2][3].

F-type ATP synthase is composed of two rotary motors termed F_o_ and F_1_. F_o_ is the membrane-embedded portion, where the *c*-oligomer ring (*c*-ring) rotates against the *ab*_2_ stator complex during proton translocation across the membrane. F_1_ is the membrane-protruding portion and rotates the inner rotor complex against the catalytic stator ring during ATP hydrolysis. F_o_ and F_1_ are connected by the rotor complex and the peripheral stalk, so as to enable the interconversion of *pmf* and the free energy of ATP hydrolysis. Under ATP-synthesizing conditions—when the *pmf* is sufficient and the rotational torque of F_o_ exceeds that of F_1_—F_o_ drives the reverse rotation of F_1_ (opposite to the ATP-hydrolyzing direction), thereby inducing ATP synthesis on the catalytic stator ring[3][4]. Conversely, when the torque of F_1_ exceeds that of F_o_ and in the absence of regulatory elements, F_1_ rotates the *c*-ring in F_o_, forcing F_o_ to pump protons in the reverse direction and thus generate *pmf*. In this way, F_o_F_1_ interconverts the *pmf* and the chemical potential of ATP hydrolysis via mechanical rotation The minimal subunit composition of F_1_, acting as the ATP-driven motor, is α_3_β_3_γ_1_, with the γ subunit embedded inside a hetero-hexameric stator ring consisting of alternating α and β subunits. The α_3_β_3_ stator ring has three catalytic sites, each located at α–β interface. Owing to the structural asymmetry of α and β, there are two distinct types of interfaces in the α_3_β_3_ ring. One-type—the α–β interface—contains the catalytic site, whereas the other type—the β–α interface—also binds ATP but does not hydrolyze it. Because most catalytically important residues reside on the β subunit, it is termed the “catalytic subunit.” Conversely, the “non-catalytic” ATP-binding site at the β–α interface is formed mainly by the α subunit residues, so the α subunit is termed the “non-catalytic subunit.” During catalysis, the three β subunits undergo conformational changes in a coordinated manner, resulting in the unidirectional rotation of the γ subunit[5].

The rotary catalysis of F_1_ has been extensively studied via single-molecule rotation assays of F_1_- ATPase derived from the thermophilic bacterium *Bacillus* PS3 (hereafter TF_1_) due to stability and the ease of handling[4]. In consistent with the pseudo-threefold symmetry of F_1_, the basic step size of rotation is 120°. This 120° step was subsequently resolved into two discrete substeps of 80° and 40°, each initiated after ATP binding and hydrolysis, respectively. Accordingly, the dwell states prior to the 80° and 40° substeps are referred to as the “binding dwell” and the “catalytic dwell,” respectively[6]. Later studies showed that F_1_ releases P_i_ from a β subunit in the catalytic dwell, but distinct from the one engaged in catalysis. It was also reported that F_1_ pauses at binding dwell angle during temperature-sensitive reaction intermediate[7]. Note that the detailed statistical analysis of the catalytic dwell revealed that ATP hydrolysis induces rotation during the dwell phase[8]. However, the angular displacement upon hydrolysis during dwell phase is subtle and within the angle distribution of the catalytic dwell (typically ±10 − 20°). Thus, TF_1_ is principally a ’6-stepper’ motor, making three binding and three catalytic dwells per turn.

Similar reaction schemes with six steps per turn have been reported for F_1_-ATPases derived from *E. coli* (EF_1_) and yeast mitochondria (yMF_1_)[9][10]. However, recent studies have revealed variations in the number of substeps. F_1_-ATPase from human and bovine mitochondrial F_1_ (*h*MF_1,_ *b*MF_1_) exhibits an additional pause—referred to as the “short dwell”—alongside the binding and catalytic dwells, resulting in nine steps per turn[11]. In contrast, F_1_-ATPase from *Paracoccus denitrificans* (PdF_1_) shows only three steps per turn under all tested conditions, despite its close evolutionary relationship to mitochondria (*Paracoccus* is an α-proteobacterium from which the mitochondrial ancestor is thought to have diverged)[12]. Thus, while most F_1_-ATPases pause six times per turn (“6-steppers”), mammalian mitochondrial F_1_ is a “9-stepper,” and PdF_1_ is a “3- stepper.” Notably, the number of steps per turn correlates with the number of the *c-*subunits in the *c*-ring: 6-steppers typically pair F_o_ with a *c*_10_-ring, 9-steppers have a *c*_8_-ring, and 3-steppers have a *c*_12_-ring. This trend suggests that F_1_ with more steps per turn has a *c*-ring containing fewer *c* subunits, implying certain mechanistic or physiological constraints on the total number of steps in the entire F_o_F_1_ complex[13].

To investigate which subunit determines the stepping pattern of F_1_, we previously constructed various hybrid F_1_-ATPases whose subunits originated from different species—TF_1_, *b*MF_1_, and PdF_1_ [14]. Analysis of these hybrids showed that rotational speed principally depends on the origin of the β subunit, as expected. However, we did not identify a single comprehensive rule governing the number of steps for all hybrids. We did find one conditional rule: whenever a hybrid F_1_ contains a subunit from PdF_1_, it consistently exhibits three-step rotation per turn, just like PdF_1_, regardless of the origins of the other subunits. Hence, although fundamental features—such as the 120° step coupled to a single ATP hydrolysis turnover and the rotation direction—are broadly conserved across species, the number and size of substeps vary. This naturally raises the question: What was the original stepping pattern of F_1_-ATPase? In other words, how did the common ancestral F_1_-ATPase rotate? Evolutionarily, F_1_-ATPase is closely related to V_1_-ATPase[15], the catalytic portion of V-type ATPases that function as ATP-driven proton pumps in the vacuoles of mammalian cells or as ATP synthases in the plasma membranes of archaea (Fig. 1A). V_1_-ATPase is also an ATP-driven rotary motor, rotating counterclockwise (when viewed from the membrane side) in the same direction as F_1_ [16]. V_1_ consists of a rotor complex and a hetero-hexameric stator ring composed of A and B subunits, corresponding to the β and the α subunits of F_1_, respectively. Like F_1_, a recent study of V_1_-ATPase from *Enterococcus hirae* (EhV_1_) resolved these 120° steps into 40° and 80° substeps [17], whereas such substeps have not been observed for *Thermus thermophilus* V_1_ (TtV_1_)[18]. Thus, although V_1_-ATPase shares some basic characteristics with F_1_-ATPase, its stepping behavior does not directly clarify how the ancestral F_1_-ATPase might have operated.

**Fig. 1.**
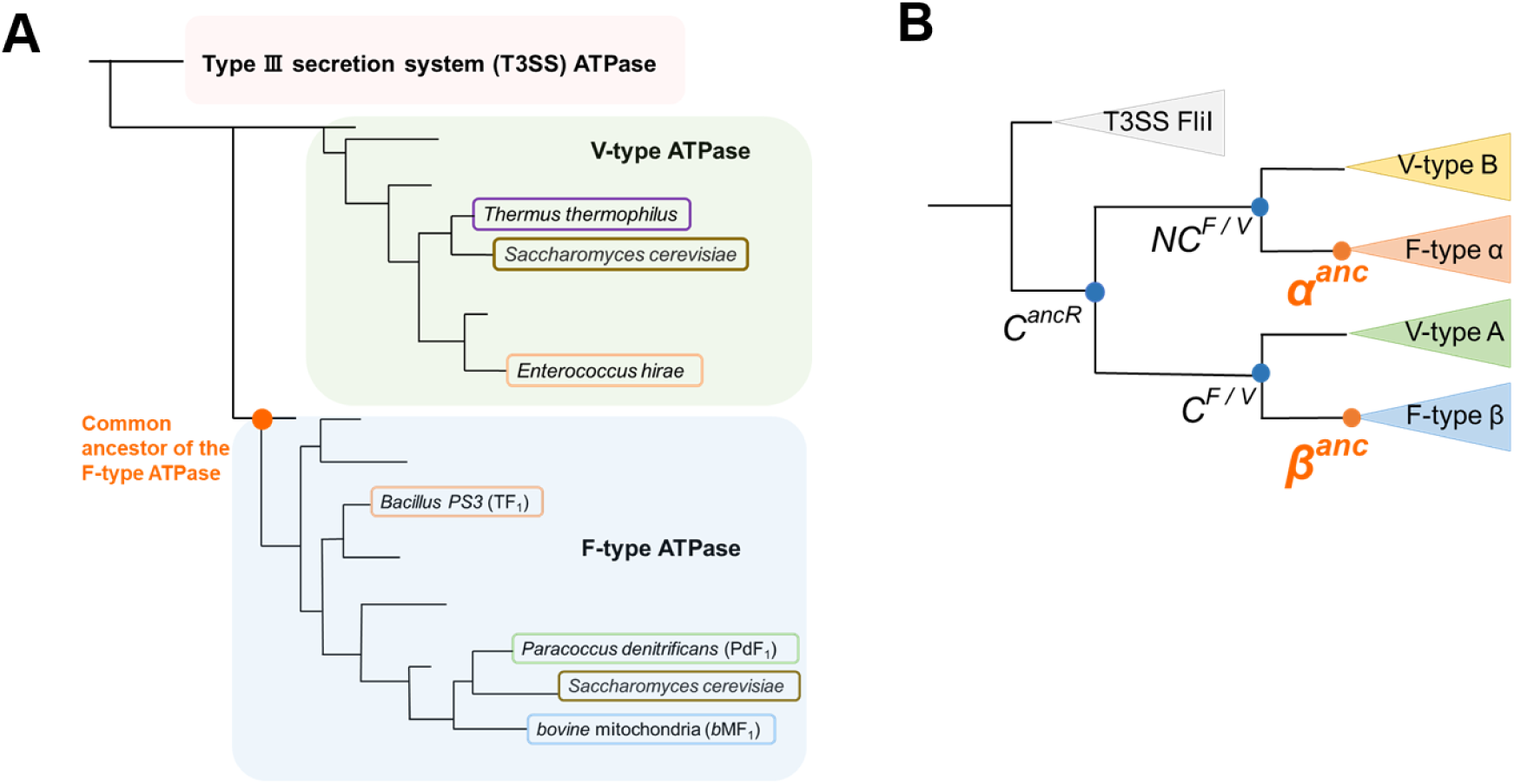
Schematic phylogenetic trees of rotary ATPases. **A.** Phylogenetic tree of rotary ATPases. The diagram highlights key species whose rotation schemes have been elucidated through single-molecule rotation assays. Species belonging to the same phylum are represented in the same color. **B.** Conceptual phylogenetic tree of the subunits forming the hexameric ring of rotary ATPases. The node labeled *C^anc^* represents the common ancestor of the α, β, A, and B subunits. *α^anc^* and *β^anc^* indicate the ancestral forms of the F-type ATPase α and β subunits, respectively, with similar annotations for other subunits. In this study, *α^anc^*and *β^anc^* are collectively considered as the ancestral form of the F-type ATPase.

One experimentally feasible way to explore the functionality of ancestral enzymes is ancestral sequence reconstruction (ASR)[19][20][21]. ASR uses multiple sequence alignment (MSA) of extant species to infer the most probable amino acid changes along a phylogenetic tree, thereby predicting ancestral protein sequences. A variety of ancestral proteins have been successfully expressed and characterized. The feasibility of sequence reconstruction by ASR methods depends on the degree of sequence conservation in the extant enzymes. Although the rotor subunits of rotary ATPases are less conserved, the non-catalytic and catalytic subunits of F_1_ and V_1_ show high sequence similarity, respectively, and even resemble each other. Molecular phylogenetic analyses suggest that the ancestral motors of F_1_-ATPase and V_1_-ATPase diverged from a common ancestral rotary motor with a hetero-hexameric stator ring comprising the ancestral non-catalytic (*NC*^F/V^) and catalytic (*C*^F/V^) subunits (Fig. 1B). It has also been proposed that this common ancestral rotary ATPase diverged from the homo-hexameric ATPase from which Type III secretion system (T3SS) ATPase originated[15][22] (Fig. 1A-B). Divergence time estimates indicate that the ancestral of either or both F_1_- and V_1_-ATPases—had already emerged as the catalytic portion of ATP synthase in LUCA[1][2].

Thus, previous studies have established an evolutionary phylogeny of rotary ATPases that stems from the pre-LUCA era[2], laying a foundation for the ancestral sequence inference of these ancient rotary ATPases. However, ancestral sequences reconstruction of multi-subunit complex enzymes requires extremely precise estimation of the amino acids forming the subunit–subunit interfaces, and consequently, there have been only a few successful examples[23]. In particular, there have been no reports of ancestral sequence research on molecular machines like F_1_-ATPase, which shows large conformational transitions.

In the present study, we reconstructed the amino acid sequences of the catalytic and non-catalytic subunits of the common ancestral F_1_-ATPase and prepared these subunits for experimental investigation to understand how the ancestral proteins functioned. We also attempted to test the functionality of the reconstituted sequences. To address the technical challenge regarding the complex instability often observed in ancestral enzymes with multi-subunit composition, we incorporated the functionally core parts from the ancestral sequences into extant thermostable F_1_, TF_1_; the non-catalytic and catalytic subunits were designed as chimeras of the ancestral and extant proteins. Specifically, the functionally critical regions—namely, the nucleotide-binding domain and the C-terminal helical domain—were derived from the ancestral sequences, whereas the N-terminal domains were taken from TF_1_ to serve as a structural scaffold. We then co-expressed these chimeric subunits with the γ subunit from TF_1_, thereby forming a functional and stable F_1_ complex, of which core parts are derived from ancestral sequence.

## Results

### Ancestral Sequence Reconstruction

Previous studies have elucidated the phylogenetic branching of subunits forming hexameric rings in T3SS ATPase, V_1_-ATPase, and F_1_-ATPase[15][22] . In this study, we focused on the five families of subunits represented in the phylogenetic tree (Fig.1B): FliI subunit comprising of the homo-hexameric ring of T3SS ATPase, the non-catalytic subunit and the catalytic subunits of the hetero-hexameric rings of V_1_- and F_1_-ATPase (the B and the A subunits for V_1_-ATPase, and the α and the β subunits for F_1_-ATPase). Representative sequence data were downloaded from National Center for Biotechnology Information (NCBI) database and used as query sequences. These query sequences were subjected to BLASTP searches[24] to collect sequences with a similarity above a certain threshold, which were then compiled into datasets for each subunit. Multiple sequence alignment was performed for each subunit. Sequences with large deletions, insertions, or redundancies were removed. As a result, we curated sequence datasets with conserved regions aligned for each subunit.

The final numbers of sequences used for phylogenetic tree construction were 94 for the T3SS ATPase FliI subunit, 128 for the A subunit and 100 for the B subunit of V_1_-ATPase, and 142 for the α subunit and 153 for the β subunit of F_1_-ATPase. Using these datasets, phylogenetic trees were inferred with IQ-TREE[25], a fast and widely used maximum-likelihood-based software that incorporates advanced model selection. The phylogenetic tree constructed was compared with those from previous research, focusing on the branching positions of phyla that were consistently present across the trees. The analysis revealed a high degree of concordance in the branching positions (Supplementary Figure 2).

The phylogenetic tree generated by IQ-TREE[25] served as a scaffold for ancestral sequence reconstruction using the codeml program in the PAML package[26], a tool for maximum-likelihood-based evolutionary analysis, and GASP[27], a probabilistic method for ancestral sequence inference. Amino acid residues at gap positions in GASP-reconstructed sequences were removed to obtain the final ancestral sequences. The reliability of the reconstructed ancestral sequences was evaluated by the probability values: approximately 0.9 for the ancestral non-catalytic subunits and catalytic subunits of F_1_-ATPase (*α^anc^* and *β^anc^*) and those for V_1_-ATPases (*B^anc^* and *A^anc^*), (Supplementary Figure 3), ensuring the high reliability of the reconstructed sequences. The common ancestral non-catalytic subunit and catalytic subunit of F_1_- and V_1_- ATPase (*NC^F/V^* and *C^F/V^*) showed the probability values around 0.65 (Supplementary Figure 3). The common ancestor protein of *NC^F/V^*and *C^F/V^*, *C^ancR^*, which is supposed to form homo- hexameric ATPase also shows a similar probability value, 0.67 (Supplementary Figure 3), suggesting the sequence reconstruction of the more upstream common ancestral proteins is less reliable. To ensure the validity of the ancestral sequence inference, we also reconstructed a phylogenetic tree using RAxML[28], a software optimized for rapid and efficient phylogenic tree searches, on which the ancestral sequence estimations was conducted. The reconstructed ancestral sequence confirms high consistency with the sequences reconstructed from the IQ-TREE-based phylogenetic trees, particularly the sequence region encoding the structurally interior parts of the subunits (Supplementary Figure 4). The sequence regions encoding subunit-subunit interfaces formed in the F_1_ complex also shows high consistency. The differences between sequences reconstructed on IQ-TREE- or RAxML-based phylogenic trees are mainly found in the structurally exterior parts and the N-terminal domains that are thought not to be crucial for functionality. Based on these results, we concluded that the ancestral sequences inferred here represent the most plausible reconstructions achievable with contemporary computational methods. In followings, we focus the sequences of *α^anc^* and *β^anc^* derived from the IQ-TREE-based phylogenic tree, while the reconstructed sequences for *NC^F/V^*, *C^F/V^* and *C^ancR^* were also analyzed (Supplementary Figure 5-8).

The sequences of the ancestral non-catalytic and catalytic subunits, *α^anc^* and *β^anc^* were compared with ones of the extant F_1_-ATPases, TF_1_, PdF_1_, and *b*MF_1_. The identities between the ancestral and the extant F_1_-ATPases were approximately 70%, while sequence identities among extant F_1_-ATPase ranged from 75% to 85% (Fig. 2A). As predicted from the high sequence conservation, we found the perfect conservation of the catalytically crucial sequences among the ancestral sequence and the extant sequences: regions including arginine finger, catalytic glutamate, and Walker motif A (phosphate binding loop, p-loop) (Fig. 2B). Thus, the sequence comparison between the sequences of the ancestral subunits and the extant ones supports the plausibility of the reconstructed sequences of *α^anc^* and *β^anc^*, suggesting the functionality of these subunits. Note that the catalytic residues are also fully conserved among *NC^F/V^*, *C^F/V^* and *C^ancR^* (Supplementary Figure 7).

**Fig. 2.**
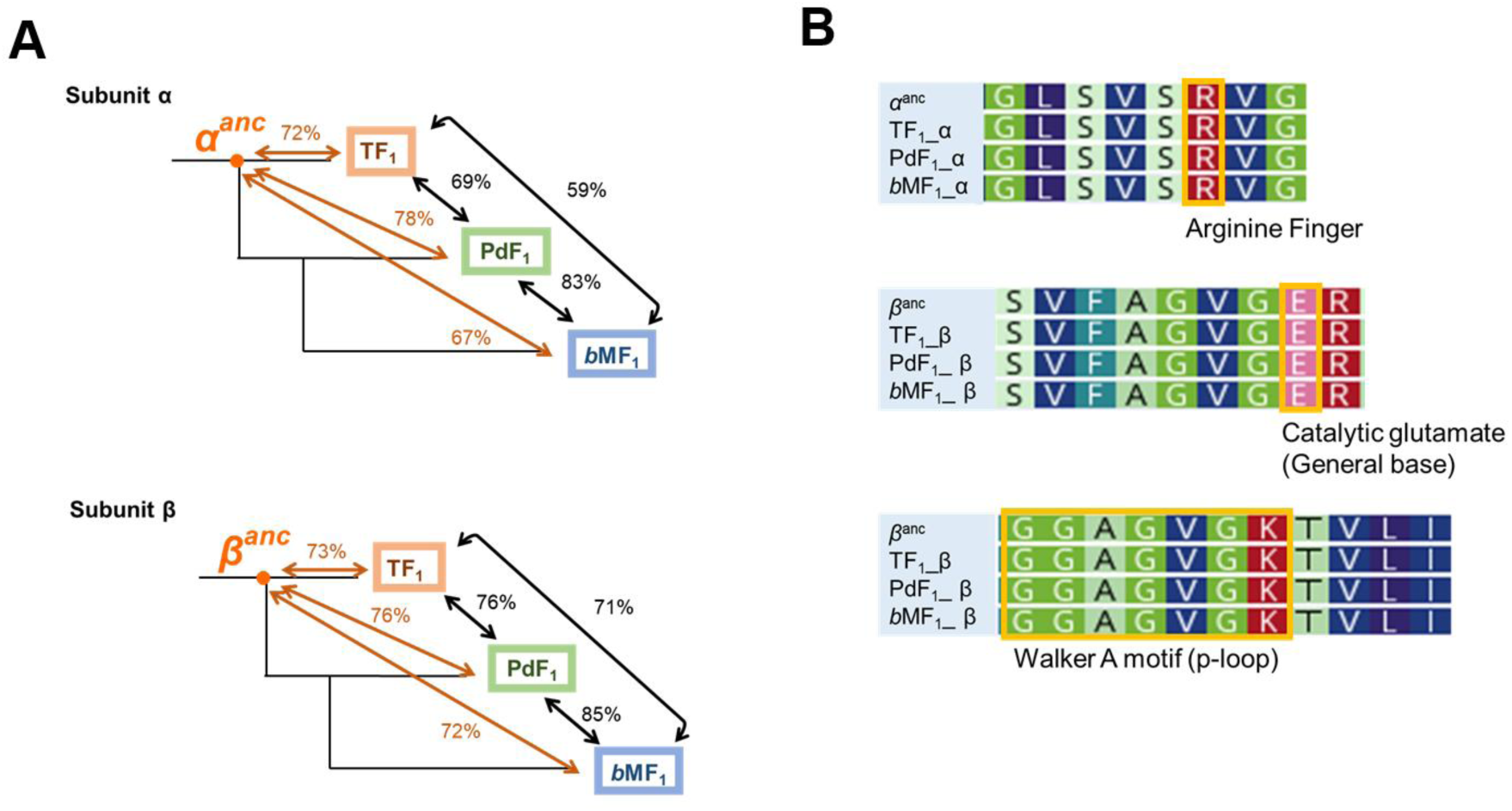
Sequence comparison between the common ancestor of F-type ATPases and extant species. **A.** Comparison of sequence identity between extant species and ancestral forms. The catalytic subunit β exhibits higher sequence identity across its full length compared to the non-catalytic subunit α. Furthermore, the sequence identity between ancestral forms and extant species shows no substantial difference compared to the sequence identity among extant species. **B.** Comparison of ATPase motifs. The conservation of key amino acid residues involved in ATP hydrolysis, including the arginine finger, catalytic glutamate, and p-loop, was analyzed. *α^anc^* and *β^anc^* represent the ancestral sequences of the F-type ATPase α and β subunits reconstructed in this study. The other three sequences correspond to the extant species TF_1_, PdF_1_, and *b*MF_1_. The ATPase motifs are well conserved across the compared sequences, suggesting that ancestral forms utilize the same key amino acid residues as extant species for ATP hydrolysis.

### Biochemical analysis

To test the functionality of the ancestral subunits, we attempted to express F_1_ composed of the ancestral subunits. ASR of enzymes with multis-subunits is challenging, due to the difficulty of highly precise inference of residues forming subunit-subunit interface. To reinforce the structural integrity, TF_1_ was used as scaffold to accommodate the functionally crucial core domains of *α^anc^* and *β^anc^*; nucleotide-binding domain (NDB) and C-terminal domain (CTD) of *α^anc^* and *β^anc^* were genetically fused with the N-terminal domain (NTD) of TF_1_ that forms the stable hexameric scaffold (residues α1-94 and β1-78 in TF_1_). The chimeric *α* and *β* subunits were co-expressed with the γ subunit of TF_1_ (Fig.3A). The resultant hybrid F_1_ with the ancestral core domains, referred hereafter to as F ^anc_core^ for simplicity, was successfully expressed as a stable complex as shown in the elution profile of size-exclusion chromatography; it showed the distinctive peak of F_1_ complexes beside peaks presumably corresponding to partially formed complexes and monomers (Fig.3B).

**Fig. 3.**
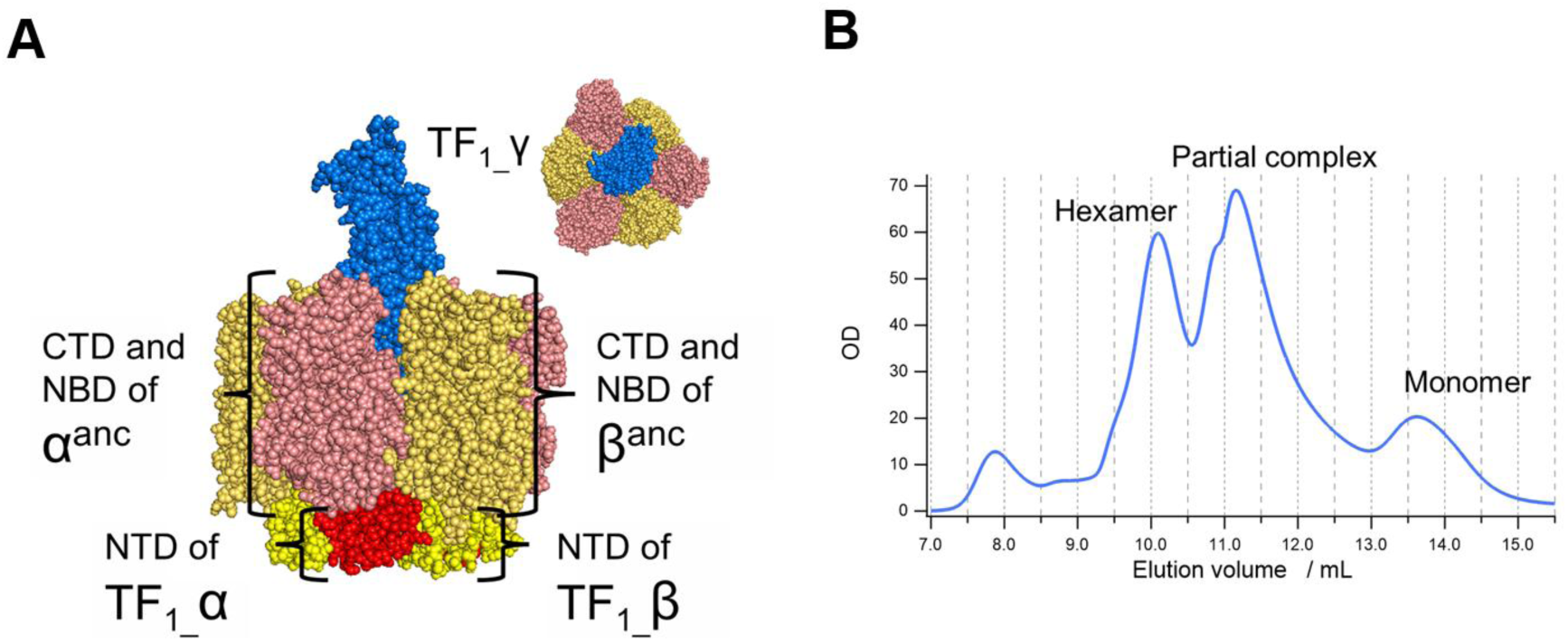
The design and purification of F_1_^anc_core^. **A.** The design of F_1_^anc_core^. The hexameric ring consists of α and β subunits, where the N-terminal β- barrel domains (NTD), forming the structural foundation, are derived from the extant species TF_1_, while the remaining regions C-terminal domain (CTD) and nucleotide-binding domain (NBD) utilize the reconstructed ancestral sequences of the F-type ATPase (*α^anc^*, *β^anc^*). The γ subunit forming the central stalk is based on the sequence of TF_1_. **B.** Size-exclusion HPLC chromatogram after Ni-NTA purification of F _anc_core_. The peak positions were estimated using a calibration curve generated with molecular weight markers. The peak corresponding to the hexameric complex aligns with the peak position observed during the purification of TF_1_, which has a comparable molecular weight. Peaks corresponding to partial complex and monomeric forms were also detected; however, the peak for the fully assembled hexameric complex is distinctly observed.

The ATPase activity of F ^anc_core^ was assessed using an ATP regeneration system. F ^anc_core^ exhibited ATPase activity approximately one-tenth that of TF_1_ (Table 1). Despite the low activity level, ATP hydrolysis was clearly evident. Notably, the ATPase activity after LDAO addition remained unchanged. LDAO is known to relieve ADP inhibition, a state in which ADP tightly remains bound on the catalytic site to prevent catalysis and rotation[29]. These findings suggest that the low ATPase activity of F ^anc_core^ reflects intrinsically low activity.

**Table 1.** ATPase activity values.

| Ancestral F<sub>1</sub>-ATPase |  | TF<sub>1</sub> |  |
| --- | --- | --- | --- |
| w/o LDAO | w/ LDAO | w/o LDAO | w/ LDAO |
| 6.7 ± 0.1 s<sup>-1</sup> | 7.5 ± 0.3 s<sup>-1</sup> | 78 ± 3 s<sup>-1</sup> | 140 ± 1 s<sup>-1</sup> |

The activity measured within the first 300 seconds of the experiment. Values are mean ± SD (n = 3).

### Structural analysis

The structure of F_1_^anc_core^ was determined by cryogenic electron microscopy (Cryo-EM), following our previous studies[30]. The purified sample was applied to EM grids at 22°C, vitrified in liquid ethane, and imaged using single-particle analysis (SPA) at 300 kV. When the grids were prepared in the presence of AMP-PNP—a nonhydrolyzable analog of ATP—only about 5% of the molecules were observed to form intact F_1_ complex, suggesting that the sample readily dissociates under those conditions (Supplementary Table 2). To overcome this, we instead prepared EM grids without any added nucleotides. Under this condition, five distinct Cryo-EM maps were obtained: two fully assembled F_1_ complexes with different γ-subunit rotational angles, a hexameric ring lacking the central shaft, and we also partially assembled complexes (a tetramer with the central shaft and a tetramer without it; see Supplementary Figure 9-11. The resolutions of the cryo-EM maps were determined to be 2.5 Å and 2.5 Å for the two fully assembled complexes, 2.8 Å for the hexameric ring without the shaft, 2.5 Å for the tetramer with the shaft, and 2.7 Å for the tetramer without the shaft, using the “gold standard” method.

A comparison of the two fully assembled complexes with previously published cryo-EM structures of TF_1_ showed a good overall match, allowing us to identify one as the catalytic dwell and the other as the binding dwell (Fig. 4A). The rotational angle difference of the γ subunit between the binding dwell and catalytic dwell in F_1_^anc_core^ was approximately 34°, smaller than that of TF_1_[30] (44°). Structural comparison of the β subunits between F_1_^anc_core^ and TF_1_ showed that they were highly similar (Fig.4B, C). Some nucleotides bound to the catalytic sites differed from those in TF_1_, likely because no exogenous nucleotides were added during sample preparation (Supplementary Fig.12-13). This suggests that the observed structures represent a state in which the enzyme re-establishes equilibrium with nucleotides originally carried over in bound form, rather than an authentic catalytic intermediate. Consistent with this overall structural similarity, the three-dimensional arrangement of conserved ATPase motif residues in the nucleotide-binding site is also well preserved between the ancestral and extant *b*MF_1_ (Supplementary Fig.14), in line with the sequence alignment shown in Fig. 2B. Although the catalytic glutamate and arginine finger are in slightly different conformations, this is likely just due to the limited resolution of the structural data.

**Fig. 4.**
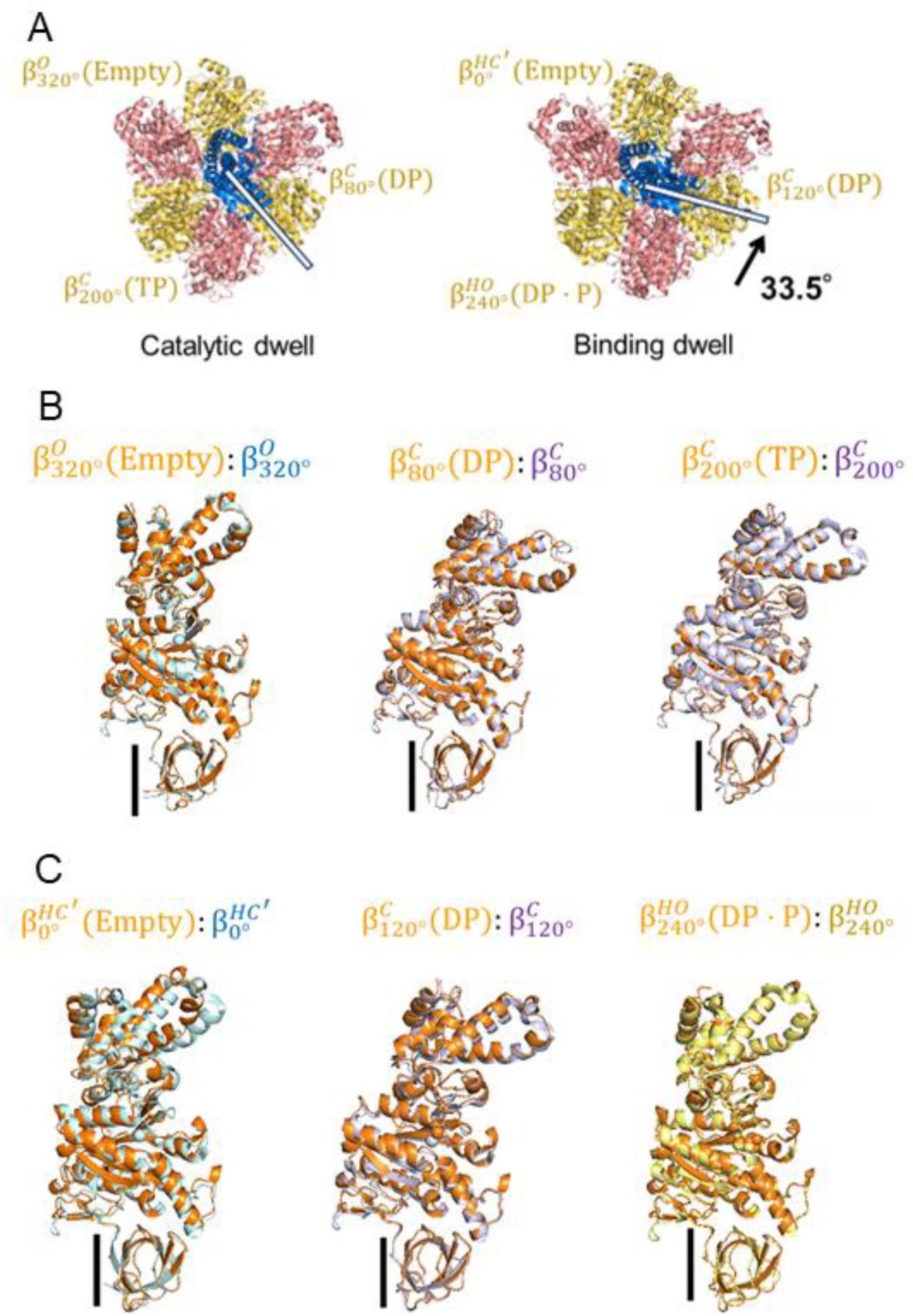
Structure of F _anc_core_. **A.** Structures of the two rotational dwells of F ^anc_core^. The structures from the top are shown, with subunits colored as in Fig. 3a. Comparison of the Catalytic dwell and Binding dwell suggests that the γ subunit rotates counterclockwise between the two dwells (rotation highlighted with white bars and a black arrow). Each dwell exhibits three distinct conformations of the β-subunits, illustrating the six sub-states (termed β^*HC*^_′_(Empty), β^*C*^ (DP), β^*C*^ (DP), β^*C*^ (TP), β^*HO*^ (DP · P), β^*O*^ (Empty)) through which the enzyme progresses during its hydrolysis cycle. **(B-C) Comparison of the β Conformations of F ^anc_core^.** The β subunit structure of F ^anc_core^ was superimposed onto the corresponding β subunit structures of TF, with a focus on the N- terminal 81 amino acid residues forming the β-barrel structure. F ^anc_core^ is depicted in orange, TF in the open conformation in blue, TF_1_ in the closed conformation in purple, and TF_1_ in the half-open conformation in yellow. **B.** Superimposition of the three β subunits in the catalytic state. The TF_1_ structure is based on PDB entry 7L1R. **C.** Superimposition of the three β subunits in the binding state. The TF_1_ structure is based on PDB entry 7L1Q.

TF_1_ is known to operate via a six-step rotational catalytic mechanism[4][30]. Given that the β subunit structures of F_1_^anc_core^ correspond one-to-one with those of TF_1_, we infer that F_1_^anc_core^ also operates via a six-step rotational catalytic mechanism similar to TF_1_. Structural alignment of the hexameric ring components (chains A–F) between F_1_^anc_core^ lacking the γ subunit and the TF_1_ binding dwell state yielded an RMSD of 0.002 Å, indicating a high degree of structural conservation within the ring. Since the hexameric ring without the central γ subunit is considered unaffected by the central γ subunit derived from TF_1_, it is highly likely that the stator ring of F_1_^anc_core^ is prone to adopt the binding dwell structure without the γ subunit. Regarding the catalytic dwell, all structural analyses of extant species to date have consistently observed the catalytic dwell structure. Taking these points into account, it is highly likely that F_1_^anc_core^ had two distinct conformational states—binding dwell state and catalytic dwell state—as same as TF_1_. However, the angular orientation of the γ subunit in the binding dwell and catalytic dwell states is influenced by the species from which the γ subunit is derived[14]. Therefore, the magnitude of the rotational angle difference between the two states in the F_1_ complex fully composed of the ancestral subunits may vary.

### Rotation analysis

Single-molecule rotation assays of F_1_^anc_core^ were performed using 40 nm gold colloid particles as the rotational probe, observed with a laser dark-field microscope at 2000 fps (frames per second)[31]. F_1_^anc_core^ rotated in a counterclockwise direction, exhibiting three distinct rotational pauses at mM level of [ATP] (Fig. 5C). Single-molecule rotation assays under various ATP concentrations allowed determination of the *V*_max_ and *K*_m_ values through Michaelis-Menten kinetics analysis, which were calculated to be 15.4 rps and 2.6 μM, respectively (Supplementary Figure 15). These *V*_max_ and *K*_m_ values were approximately one-tenth of those observed for TF_1_. The rate constant of ATP binding, *k*_on_, estimated as 3×*V*_max_ / *K*_m_, was determined to be 1.6×10^7^ M^-1^s^-1^, which is very close to that of TF_1_ (1.8×10^7^ M^-1^s^-1^). Thus, the major kinetic difference is the slow maximum rotation speed.

**Fig. 5.**
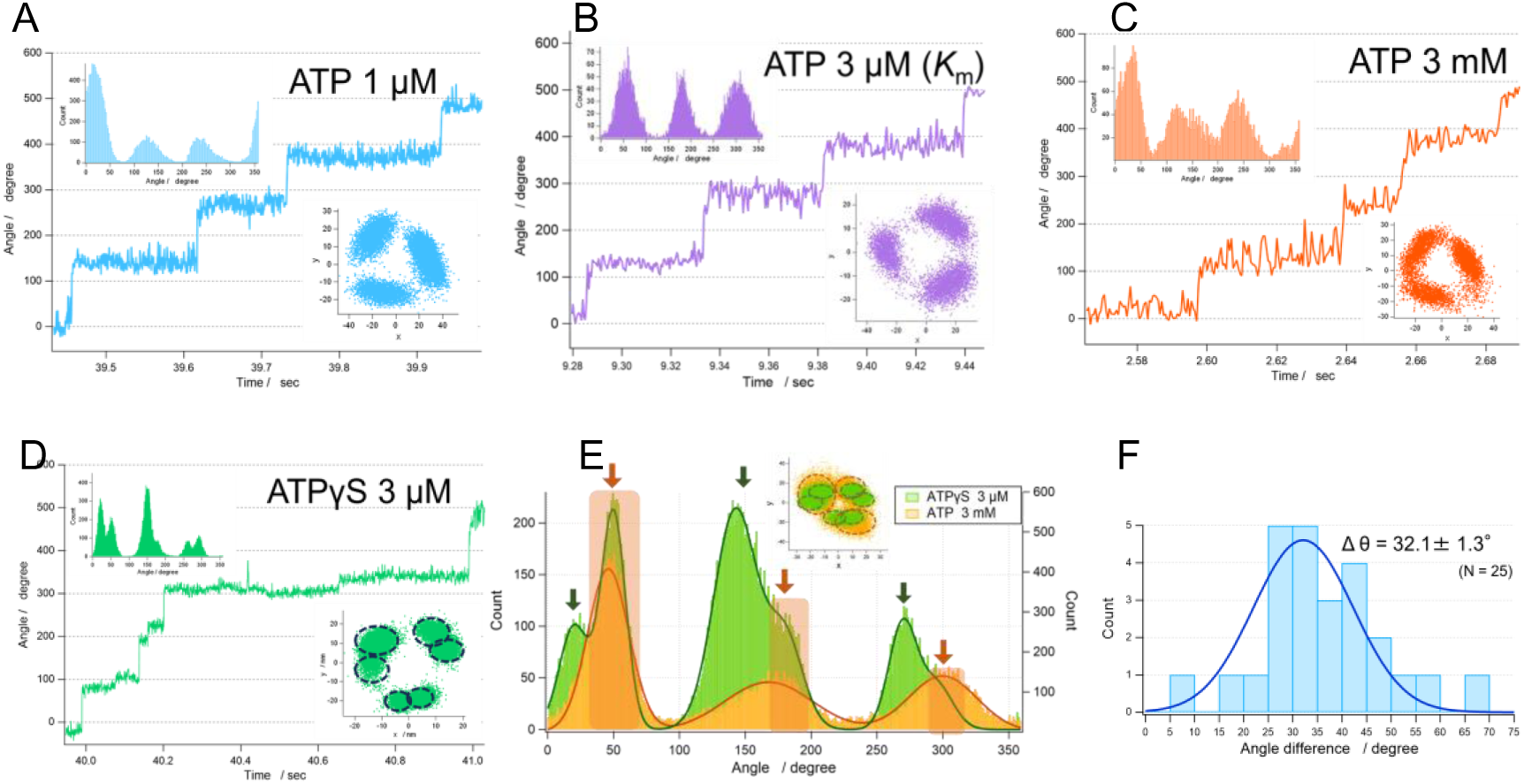
Single-molecule rotation assay of F_1_^anc_core^. (A-D) Time courses of rotation. The angular position histograms (left), and the x-y plots of the centroid of a rotating gold colloid (right) are shown in inset. **A.** 1 μM ATP **B.** 3 μM ATP (*K*_m_). **C.** 3mM ATP. **D.** 3 μM ATPγS. **E.** The angular position histograms of 3 mM ATP (orange) and 3 μM ATPγS (green). The x-y plots are shown in inset. The arrows show the angular position of catalytic dwell (green) and binding dwell (orange). **F.** Histogram of angular differences between catalytic dwell and binding dwell positions. Values are mean ± SD (n = 25, 10 molecules).

The maximum rotational velocity 15.4 rps corresponds to 46/sec as *k*_cat_ of hydrolysis, that is significantly faster than the rate of hydrolysis estimated from biochemical ATPase activity assays performed with ATP regeneration system (Table 1). Such a discrepancy between the single-molecule rotation assay and the biochemical assay has often been reported[11][12][32], and it is usually attributed to ADP-inhibition. In the presence case, in addition to ADP-inhibition, the apparently lower hydrolysis activity estimated from biochemical analysis can be attributed to the heterogeneity of the sample. Cryo-EM analysis revealed that a significant fraction of the sample consisted of partial complexes lacking the γ subunit and αβ pair.

Another distinctive feature of F_1_^anc_core^ rotation is that it proceeds in three steps per turn at all of [ATP] examined (Fig. 5A-C), whereas TF_1_ is known to make 80° and 40° substeps during each 120° rotation, particularly around *K*_m_ region making six steps per turn. The buffer exchange experiment where [ATP] was switched between high and low concentrations confirmed that the dwell positions observed at low and high [ATP]s were coincident with each other (Supplementary Figure 16). This observation can be interpreted to mean that F_1_^anc_core^ performs both ATP binding and hydrolysis within the same dwell. However, such an interpretation contradicts the cryo-EM analysis, which shows that F_1_^anc_core^ adopts two distinct conformational states—a binding dwell and a catalytic dwell—where the γ subunit’s orientation differs by 33.5° (Fig. 4A). Another possibility is that F_1_^anc_core^ pauses for a prolonged period at the binding dwell angles while waiting on a reaction step other than ATP binding, and that the long pause dominates the overall reaction time, making the shorter catalytic dwells effectively negligible.

To test this hypothesis, we conducted the rotation assay in the presence of ATPγS, a slowly hydrolyzed ATP analog often used to identify the angular position of the catalytic dwell in the rotation assay of F_1_[11][12][33]. As expected, F_1_^anc_core^ rotated at significantly lower speed in ATPγS than in ATP; the maximum rate was 1.46 rps, which is around one-tenth of ATP-driven rotation (Supplementary Figure 17). When the rotation was observed in the presence of 3 μM ATPγS— near the *K*_m_ range for ATPγS-driven rotation (0.68 μM)—six rotational dwells were identified, separated by 32.1°, consistent with the predictions from cryo-EM analysis. To correlate these ATPγS-based dwell positions with those observed under ATP, we performed solution exchange experiments, alternately supplying ATP and ATPγS. By comparing the dwell positions in both conditions, we found that three of the six dwell points observed with ATPγS matched those under ATP. Based on this result, we assigned the three shared dwell points, seen with both ATP and ATPγS, to the binding dwell, whereas the dwell points observed exclusively under ATPγS were identified as the catalytic dwells. The analysis of the angular differences of the binding dwell from the catalytic dwell revealed a histogram centered at 32.1°. This angular difference aligns closely with the rotational angle difference of ∼34° between the binding and catalytic dwells of the γ subunit observed in Cryo-EM structural analyses, supporting the above assignment. Thus, it was confirmed that F ^anc_core^ has two distinct states pausing at binding dwell angles or catalytic dwell angles as expected from Cryo-EM analysis, while the motor pauses predominantly at binding dwell angles waiting for a long reaction step to occur under ATP conditions. TF_1_ is reported to make pauses at binding dwell angles when observed at low temperatures, due to temperature sensitive reaction (TS reaction). To test the possibility of TS reaction, we measured ATP hydrolysis activity at low temperatures. However, *Q*_10_ factor estimated from ATPase assay for F ^anc_core^ was 1.52, too low to attribute the rate-limiting step as TS reaction (Supplementary Table 4). Thus, the long pause found at binding dwell angles should be a new class of reaction or conformational state.

## Discussion

In this study, we focused on the phylogenetic tree of rotary ATPases and inferred ancestral sequences of the hexameric ring–forming subunits at multiple nodes: the ancestral F_1_-ATPase, the common ancestral rotary ATPase of F_1_- and V_1_-ATPases, and an even earlier rotary ATPase with a homo-hexameric ring. As a result, we obtained subunit sequences for these ancestral rings—*α^anc^*, *β^anc^*, *C^F/V^, NC^F/V^* and *C^ancR^*. Comparative analysis revealed that these ancestral rotary ATPases already possessed the key motifs required for ATP hydrolysis—namely, the arginine-finger, catalytic glutamate, and Walker motif A—that are almost universally found in extant F_1_- ATPases. (Fig. 2B, Supplementary Figure 7). Although we did not perform a quantitative structural analysis, the predicted structures of these ancestral subunits closely resemble those of modern subunits (Supplementary Figure 8).

Considering that LUCA employed either the ancestral rotary ATPase corresponding to F_1_ or V_1_-ATPase (or both)[1][2], the common ancestor of F_1_ and V_1_ must have already arisen as a rotary ATPase with a hexameric ring in the pre-LUCA era. It is also plausible that the ancestral ATPase diverging from the Type III secretion system (T3SS) ATPase possessed a hexameric ring structure, given that extant T3SS ATPases likewise form hexamers[34] [22].

Before expressing F ^anc_core^, which carries *α^anc^* and *β^anc^* sequences in its core domains, we first attempted to produce several ancestral ATPases—namely *C^F/V^*, *NC^F/V^*, and *C^ancR^*. However, none of these proteins formed stable complexes, indicating that it remains challenging to obtain functional complexes solely from ancestral sequences with our current knowledge of rotary ATPase and the present ASR methodology. To address this issue, we constructed a chimeric ATPase using an extant F_1_ (TF_1_) as a structural scaffold. Specifically, we combined the TF_1_ N-terminal domain (NTD), which forms the base of the hexameric ring, with the ancestral sequences for the functionally essential domains—the nucleotide-binding domain (NBD) and the C-terminal domain (CTD) of *α^anc^* and *β^anc^*. When expressed with the γ subunit of TF_1_, this design yielded a stable F_1_ complex, termed F ^anc_core^.

F_1_^anc_core^ was expressed as a stable complex exhibiting ATPase activity although the activity is significantly lower than TF_1_. Cryo-EM structural analysis revealed the significant fraction of molecules retained the α_3_β_3_γ complex although partial complexes such as α_3_β_3,_ α_2_β_2_γ, or α_2_β_2_ subcomplexes were also found. The α_3_β_3_γ complex structure of F ^anc_core^ exhibited the two conformations. Comparison with TF_1_ structures identified the two conformational states as the binding dwell state and catalytic dwell state, indicating that F ^anc_core^ shares the same rotational catalytic mechanism as same as TF_1_. Interestingly, the conformational states of the α_3_β_3_ subcomplex corresponds to the binding dwell state as found in the α_3_β_3_ subcomplex of TF_1_, reinforcing the abovementioned idea that F ^anc_core^ operates the similar rotational catalytic mechanism to TF_1_, alternating the conformational state between binding dwell state and catalytic dwell.

The result of rotation assay of F ^anc_core^ was apparently against this expectation. F ^anc_cor^ showed a 3-step rotation at all tested [ATP], indicating that it consistently pauses at binding dwell angles. Presuming that this apparent 3-step pattern arises from an extended binding dwell overshadowing a brief catalytic dwell, we conducted rotation assays with ATPγS. Under these conditions, we observed well-defined stepping, with six pauses per turn. A subsequent buffer-exchange experiment—switching between ATP and ATPγS—revealed that F ^anc_core^ pauses at both binding and catalytic dwell angles, which differ by 32.1°. This value closely matches the ∼34° difference determined by Cryo-EM analysis.

Thus, F ^anc_core^ essentially follows a reaction scheme similar to that of TF_1_, featuring two stable conformational states (binding dwell and catalytic dwell), even though the binding dwell of F ^anc_core^ is significantly longer than its catalytic dwell (Supplementary Figure 18). The molecular mechanism behind this prolonged binding dwell remains unclear. We hypothesized it might stem from a temperature-sensitive process, akin to what has been observed in TF_1_ rotation assays at low temperatures. However, the *Q*_10_ factor for the ATPase activity of F ^anc_core^ is only 1.52—too low for a typical temperature-sensitive reaction. A candidate reaction responsible for the long-lived dwell is ADP release step that was suggested at or near the binding dwell angles[6][35]. Further investigation remains to address this point.

Extant F_1_-ATPases exhibit diverse rotary mechanisms, classified as 3-steppers, 6-steppers, or 9-steppers. Our findings suggest that the ancestral F_1_-ATPase was fundamentally a 6-stepper. One might argue that if the binding dwell is relatively longer than the catalytic dwell, the motor could effectively appear to be a 3-stepper. Yet, PdF_1_, the only known 3-stepper motor, still displays 3-step rotation even when measured with ATPγS, differing it from F_1_^anc_core^ described here. Consequently, it seems more appropriate to conclude that the ancestral F_1_ was intrinsically a 6- stepper, and it diversified into more 3-steppers, 6-steppers, and 9-steppers (Fig. 6).

**Fig. 6.**
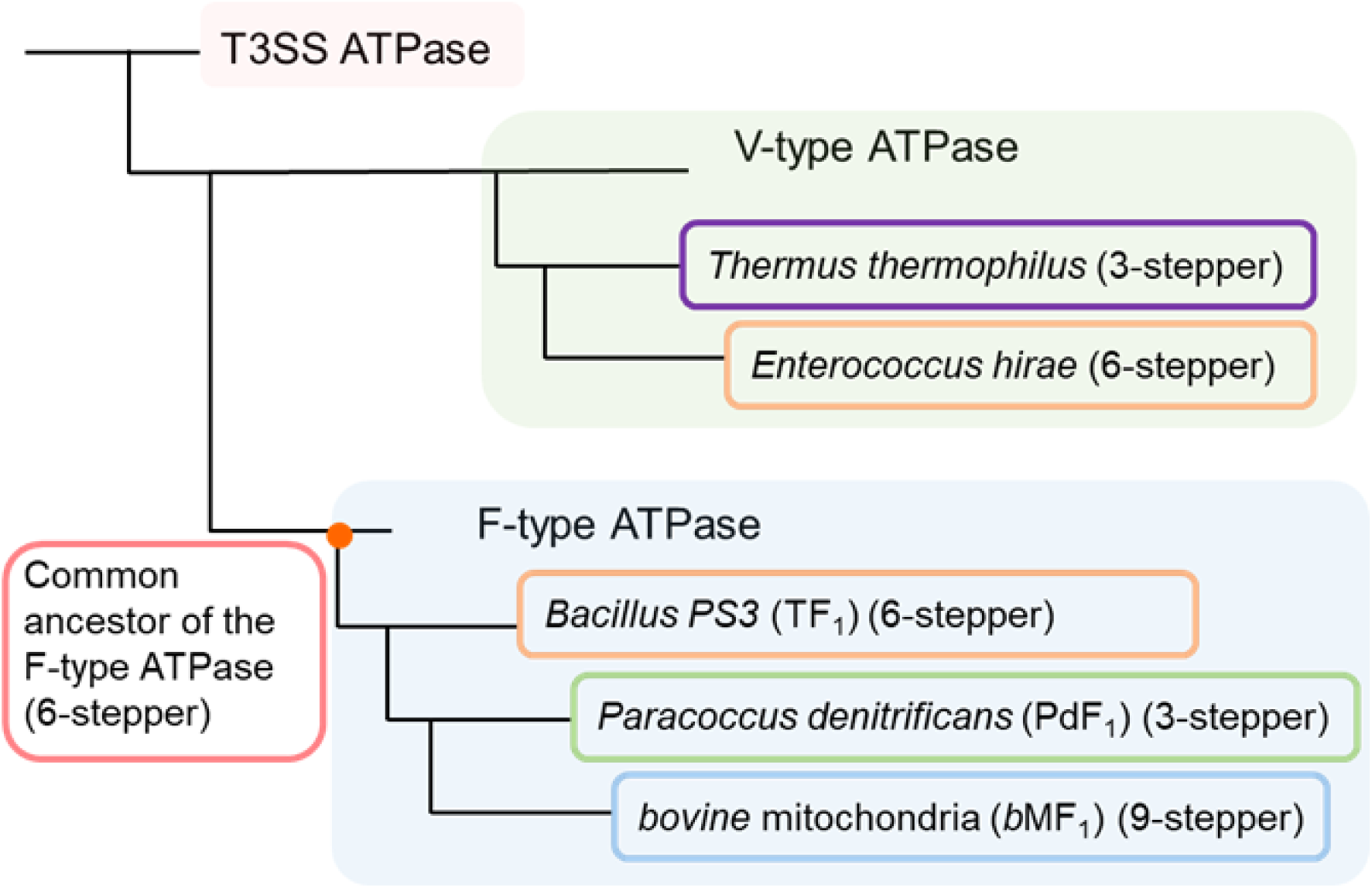
Differences in Rotational Catalytic Mechanisms Among Species and Their Branching Positions. The branching positions and unique features of the rotational catalytic mechanisms for species with well-characterized mechanisms are presented.

We previously identified an intriguing correlation between the number of rotational steps of F_1_ and the number of the *c*-subunits in the *c*-ring[13], although more data is needed to confirm its generality. From this correlation, one empirical rule emerges: a 6-stepper F_1_ pair with a 10-stepper F_o_ (i.e., 10 *c*-subunits). This rule suggests that ancestral F_1_ likely paired with a F_o_ containing a *c*_10_- ring. If the free energy of ATP hydrolysis was comparable to that of extant cells, then LUCA and related lineages must have already possessed electrically tight plasma membranes capable of maintaining a sufficiently high *pmf*. This follows from the fact that the number of *c*-subunits is one of the key factors determining the H⁺/ATP ratio and the equilibrium *pmf* between ATP hydrolysis and synthesis.

From an engineering perspective, this study highlights a new strategy: using TF_1_ as a scaffold for implementing various ancestral or designed catalytic domains, including the common subunits of F_1_- and V_1_-ATPase, *C*^F/V^and *NC*^F/V^. By extending this idea to the entire F_o_F_1_ complex, one could consider incorporating more ancestral-sequence subunits into a stably structured F_o_F_1_—such as T F_o_F_1_ found in extant species—to more accurately estimate the function of ancestral F_o_F_1_. Currently, ancestral sequence reconstruction methods are largely restricted to highly conserved subunits, so beyond the α and β subunits of F_1_, they can only be applied to certain regions of the *c*-subunit or the *a*-subunit. Even so, studying chimeric or hybrid F_o_F_1_ complexes that include such ancestral functional units may yield insights into functions and dynamics that cannot be inferred from sequence analysis alone.

## Methods

### Phylogenetic analysis and ancestral sequence inference

Protein sequences were retrieved from the NCBI (National Center for Biotechnology Information) database. In this study, the KF database ver.2021[36] was further expanded for ATPase research, resulting in the KFS Database. The database comprised all protein sequences of 136 archaeal species, 594 bacterial species, and 77 eukaryotic species, and was employed for BLAST searches.

Amino acid sequences for the α and β subunits of the F_1_-ATPase from *bovine* mitochondria (ACCESSION: P19483, P00829), the A and B subunits of the V_1_-ATPase from *Thermus thermophilus* (Q56403, Q56404), and the T3SS FliI protein from *Escherichia coli* (NP_416451) were retrieved from NCBI as query sequences. The FliI family of the T3SS was incorporated into the phylogenetic tree as an outgroup to determine the root position of F_1_-ATPase and V_1_-ATPase. These sequences were subjected to BLASTP searches[24] against the KFS Database to collect homologous sequences. Redundant sequences and those with significantly different lengths were removed. The remaining 617 sequences were realigned using MAFFT[37] with secondary structure considerations, followed by manual refinement to produce a multiple sequence alignment, which was subsequently used for phylogenetic analysis.

Alignment trimming was performed using TrimA_l_[38] (version 1.4) in automated trimming mode (- automated1). Phylogenetic trees were constructed using IQ-TREE[25] (version 2.1.3). The ModelFinder Plus[39] (MFP) module identified the LG+R10 model as the optimal amino acid substitution model. Ultra fast bootstrap analysis[40] with 1,000 replicates was conducted to evaluate branch confidence.

For additional reliability assessments, phylogenetic analysis was also conducted using RAxML[28] (version 8.2.12). The LG+G+I model was selected, and bootstrap analysis with 1,000 replicates was performed.

Phylogenetic analyses and ancestral sequence reconstruction were performed using the codeml program in the PAML package (version 4.9j, February 2020) [26] with the LG substitution model for amino acid sequences. Default parameters were used unless specified otherwise. The gap positions of ancestral sequences were inferred by GASP[27] (version 2.0.0).

The ancestral sequences generated by PAML were compared with those by GASP. Regions identified as gaps in the ancestral sequence of GASP were excised from the PAML ancestral sequences to produce the final ancestral sequence set.

### Preparation for F _anc_core_

Genuine TF_1_ was prepared as described[41]. To visualize rotation, two cysteine residues were introduced as previously reported[14]. The plasmid encoding F ^anc_core^ was constructed using the TF_1_ plasmid as a vector and PCR products encoding the C-terminal and nucleotide-binding domains of *α*_anc_ and *β*_anc_ (both with N-terminal His-tags) as insert DNAs. After ligation, the recombinant plasmid was introduced into the F_o_F_1_-deficient *E. coli* strain JM103Δunc. F ^anc_core^ was then expressed in *E. coli*, purified, and biotinylated as described in [42]. The protein concentration was determined by UV absorbance using a molar extinction coefficient of 182,500 M⁻¹ cm⁻¹, calculated from its amino acid sequence using the ProtParam tool (ExPASy).

### Calibration Curve for Molecular Weight Determination by Size-Exclusion HPLC

Molecular weight determination was performed using size-exclusion high-performance liquid chromatography (SEC-HPLC). A calibration curve was generated using the MW-Marker (HPLC) for Molecular Weight Determination (Oriental Yeast Co., Ltd.). Chromatographic analysis was conducted on an HPLC system equipped with a size-exclusion column maintained at 25°C. The mobile phase consisted of 50 mM HEPES-KOH (pH 7.5) containing 100 mM NaCl, delivered at a flow rate of 0.5 mL/min. UV absorbance was monitored at 280 nm.

Retention times of the marker proteins were recorded, and a calibration curve was constructed by plotting the logarithm of molecular weight against retention time. A linear regression analysis was performed to establish the calibration equation, which was subsequently used to estimate the molecular weight of F_1_^anc_core^. Calibration was validated through triplicate measurements, ensuring reproducibility within an acceptable standard deviation.

### ATPase activity assay

ATPase activity was measured at 25°C in a buffer containing 50 mM Hepes-KOH (pH 7.0), 50 mM KCl, 3 mM MgCl₂, and an ATP-regenerating system (0.2 mM NADH, 2.5 mM phosphoenolpyruvate, 200 μg/mL pyruvate kinase, and 50 μg/mL lactate dehydrogenase). Activity was calculated from the slope of NADH absorbance during the first 300 seconds of measurement. LDAO-stimulated activity was assessed by adding 0.3% LDAO to the reaction and calculating the slope of NADH absorbance thereafter.

### Cryo-EM grid preparation

A volume of 3.5 μL of purified F ^anc_core^ was applied to a glow-discharged holey gold grid (Ultrafoils R0.6/1.0, 200 mesh). The grids were blotted for 4 seconds at 22°C and 100% humidity, then plunge-frozen in liquid ethane using a FEI Vitrobot Mark IV.

### Data collection

The grids were initially screened for ice thickness and particle density using a Thermo Fisher Scientific Talos Arctica transmission electron microscope (TEM) operating at 200 kV. Subsequently, the grids were transferred to a Thermo Fisher Scientific Titan Krios TEM operating at 300 kV, equipped with a Gatan BioQuantum energy filter (20 eV slit width) and a K3 camera. To mitigate orientation bias observed in an initial test sample, movie micrographs were recorded at tilt angles ranging from 20° to 40°, as suggested by cryoEF[43], which indicated an optimal tilt angle of ∼37°. Automatic data collection was performed using EPU (E Pluribus Unum, Thermo Fisher Scientific) at a nominal magnification of ×60,000 (displayed magnification of ×165,000 due to the energy filter), resulting in a pixel size of 0.84 Å. The total electron dose was set to 62 electrons per Å², distributed over 80 frames with a total exposure time of 6.2 seconds. A total of 4373 movie micrographs were collected.

### Data processing

All image processing and refinement were performed using CryoSPARC v4.4.1[44]. Initially, micrographs were motion-corrected, and defocus values were estimated using the patch-based workflow. Particles were automatically picked and subjected to two-dimensional (2D) classification to exclude “junk” particles, such as those from aggregates or minor contaminants. The 2D classes were split into hexamers and tetramers and processed separately. Initial models were generated by *ab initio* classification into 2 -3 classes. These *ab initio* maps were then used as inputs for heterogeneous refinement and 3D classifications, allowing further classification of particles into distinct structural classes. Each class was independently processed through homogeneous refinement and non-uniform refinements, yielding the final high-resolution maps. In regions where lower-resolution features were observed, DeepEMhancer[45] was applied to sharpen the maps, enhancing their interpretability in figures displaying the entire complex.

### Model building

Models were constructed for intact F_1_ complexes and refined using Coot[46] and PHENIX[47], with PDB structures 7L1Q (TF_1_ binding dwell cryo-EM structure) and 7L1R (TF_1_ catalytic dwell cryo-EM structure) serving as templates. Supplementary Table 3. shows details of the refinement and validation statistics. Figures made using PyMOL.

### Single-molecule rotation assay

Flow chambers were constructed using double-sided tape as spacers and two cover glasses (18 × 18 mm² and 24 × 32 mm²; Matsunami Glass). The bottom glass surface was coated with Ni-NTA.

The basic assay buffer contained 50 mM Hepes-KOH (pH 7.5), 100 mM KCl, and 5 mM MgCl₂. When ATP was used as the substrate, an ATP-regenerating system (2 mM phosphoenolpyruvate and 100 μL/mL pyruvate kinase) was added.

The flow chamber was first incubated with basic buffer containing 5 mg/mL BSA (BSA buffer) for 5 minutes. F_1_ molecules (200–500 pM) in BSA buffer were then introduced and incubated for 10 minutes, followed by washing with BSA buffer to remove unbound molecules. Next, 40 nm gold nanoparticles (prepared as described in [17]) were introduced, incubated for 10 minutes, and unbound particles were removed by washing with substrate-containing basic buffer.

Rotational assays were performed as described in [17] at room temperature. Recorded videos were analyzed using custom software.

### Data availability

The cryo-EM maps from this study are deposited in the Electron Microscopy Data Bank (EMDB) under accession codes 49841, 49839, 49840, 49843 and 49842 for hexamer without stalk, hexamer with stalk in binding dwell, hexamer with stalk in catalytic dwell, tetramer without stalk, and tetramer with stalk, respectively. The models generated and analyzed for the two of the above maps are available in the Protein Data Bank under accession codes 9NVL for hexamer with stalk in binding dwell, and 9NVM for hexamer with stalk in catalytic dwell.

