## Supporting Information for "Functional and Structural Characterization of F_1_-ATPase with common ancestral core domains in stator ring"

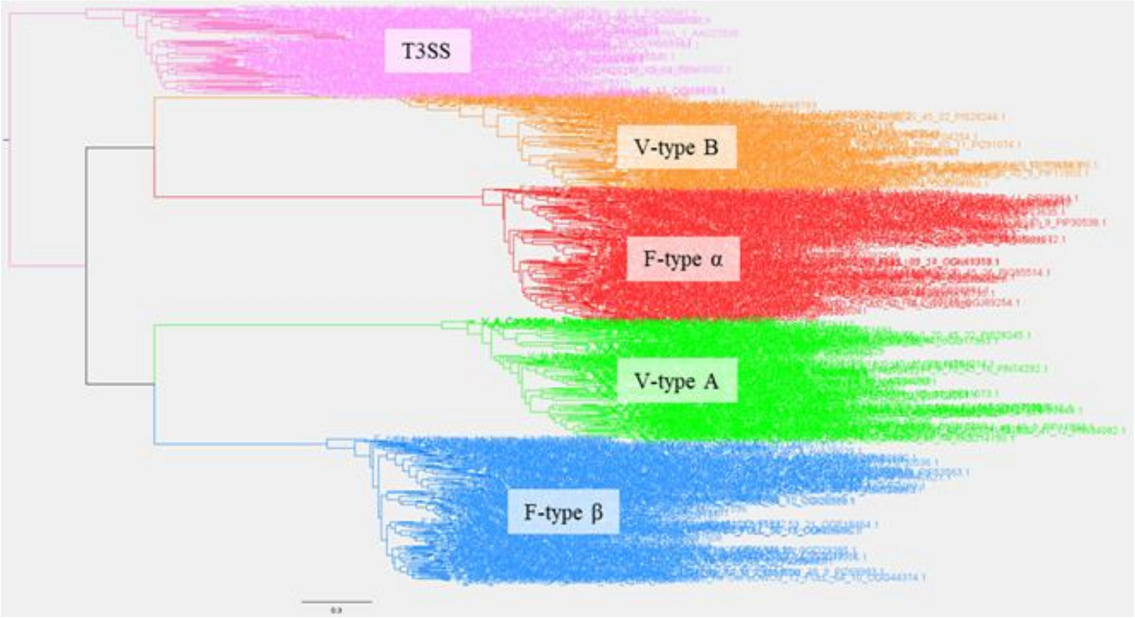


**Fig. S1. Phylogenetic tree of catalytic/non-catalytic subunits of rotary ATPases**

The phylogenetic tree was constructed using IQ-TREE based on a dataset of 617 sequences, including 94 T3SS ATPase FliI sequences, 142 F-type ATPase α-subunit sequences, 153 β-subunit sequences, 128 V-type ATPase A-subunit sequences, and 100 B-subunit sequences.

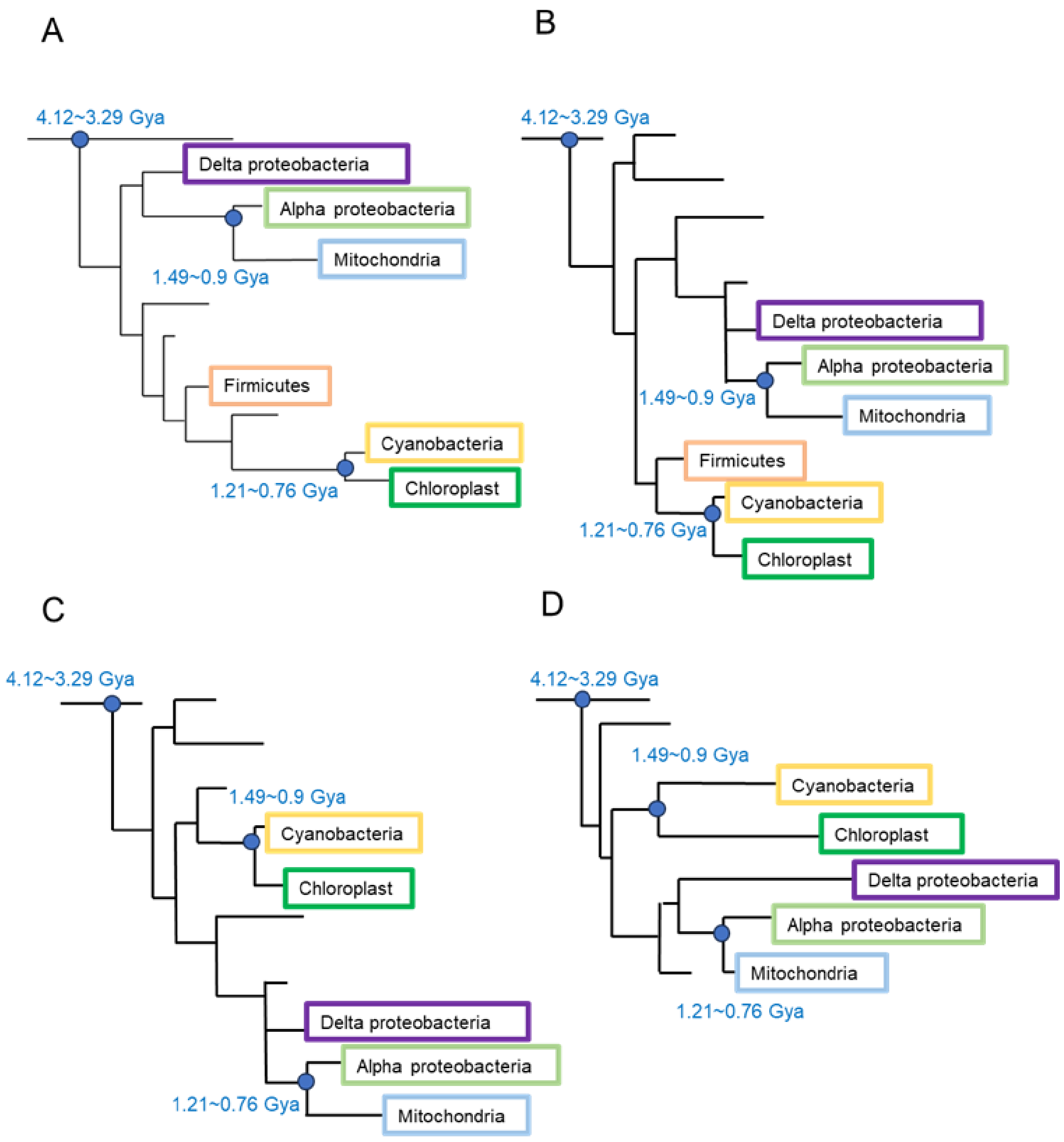


**Fig. S2. Phylogenetic Tree Topology Comparison**

To compare tree topologies, we manually extracted the branching patterns of shared phyla. Branch lengths indicate evolutionary rates. Estimated divergence times are annotated at corresponding nodes based on the time estimates from a previous study[1]. **(A-B)** α subunit. **A.** Previous study[1], **B.** This study. **(C-D)** β subunit. **C.** Previous study[1], **D.** This study.

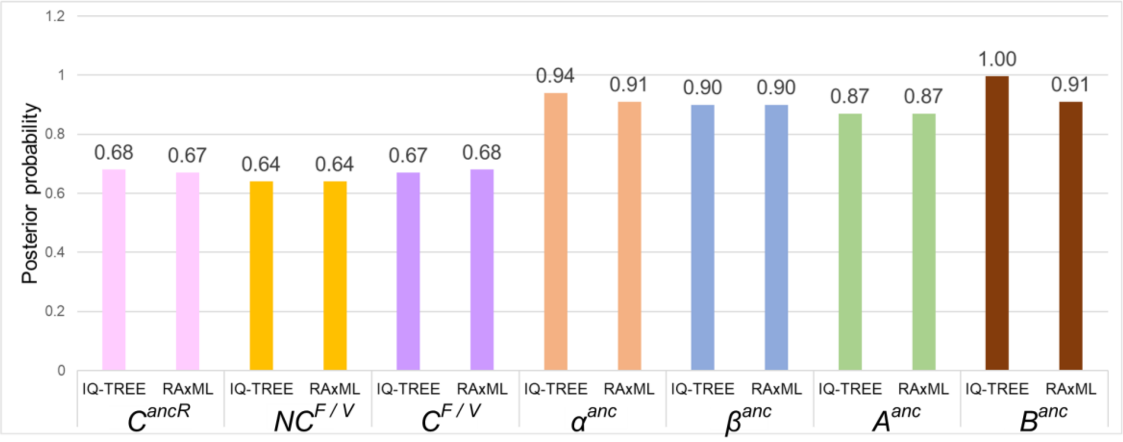


**Fig. S3. Evaluation of Posterior Probability Values**

Posterior probability values of ancestral sequences reconstructed with IQ-tree-based or RAxML-based phylogenetic trees are presented for each ancestral subunit.

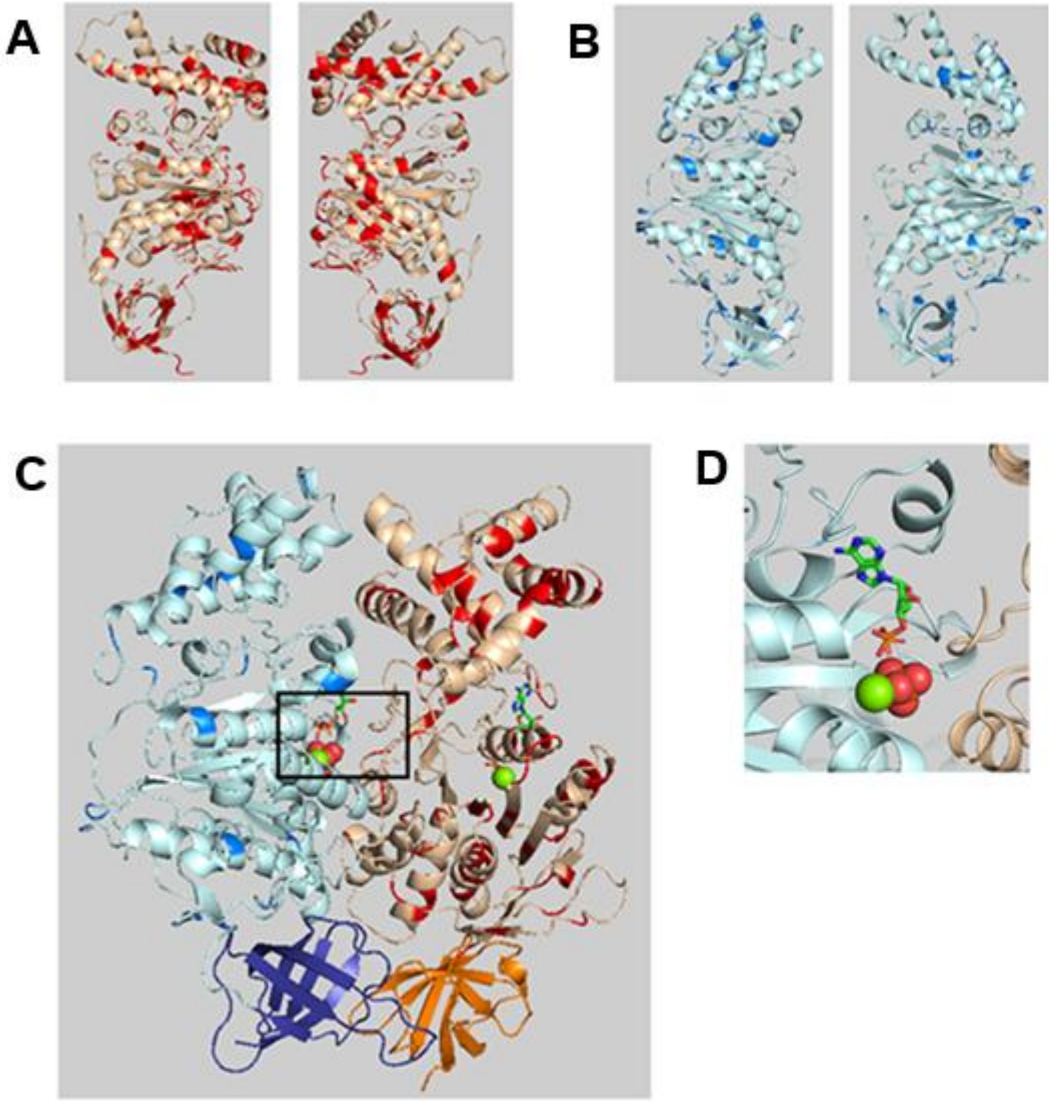


**Fig. S4. Comparison of IQ-TREE- and RAxML-based sequence reconstruction**

The sequences of the ancestral subunit inferred with IQ-TREE- and RAxML-based phylogenic trees were compared and the sequence differences were mapped on the subunit structure of $TF_1$ The structural information was obtained from the PDB (PDB ID:4XD7). **A.** Structural map of the common ancestral α-subunit ($\alpha^{anc}$). The sequence differences are highlighted in red, while conserved regions are shown in cream. **B.** Structural map of the common ancestral β-subunit ($\beta^{anc}$). The sequence differences are highlighted in red, while conserved regions are shown in cream. **C.** Structural map of the α-β interface. The N-terminal domains of the α- or β-subunit are depicted with dark blue and orange. **D.** A magnified view of the substrate-binding site. No amino acid difference was observed between the two sequences.

**Table. S1. Sequence Identity Between Ancestral and Extant sequences**

| | $C^{ancR}$ | | $NC^{F/V}$ | | $C^{F/V}$ | |
|---|---|---|---|---|---|---|
| | IQ-TREE | RAxML | IQ-TREE | RAxML | IQ-TREE | RAxML |
| $TF_1$_α | 37 % | 35 % | 43 % | 39 % | | |
| $TF_1$_β | 46 % | 46 % | | | 58 % | 56 % |
| *b*$MF_1$_α | 36 % | 35 % | 43 % | 40 % | | |
| *b*$MF_1$_β | 43 % | 45 % | | | 58 % | 56 % |
| $TtV_1$_A | 34 % | 33 % | | | 40 % | 40 % |
| $TtV_1$_B | 36 % | 37 % | 39 % | 41 % | | |

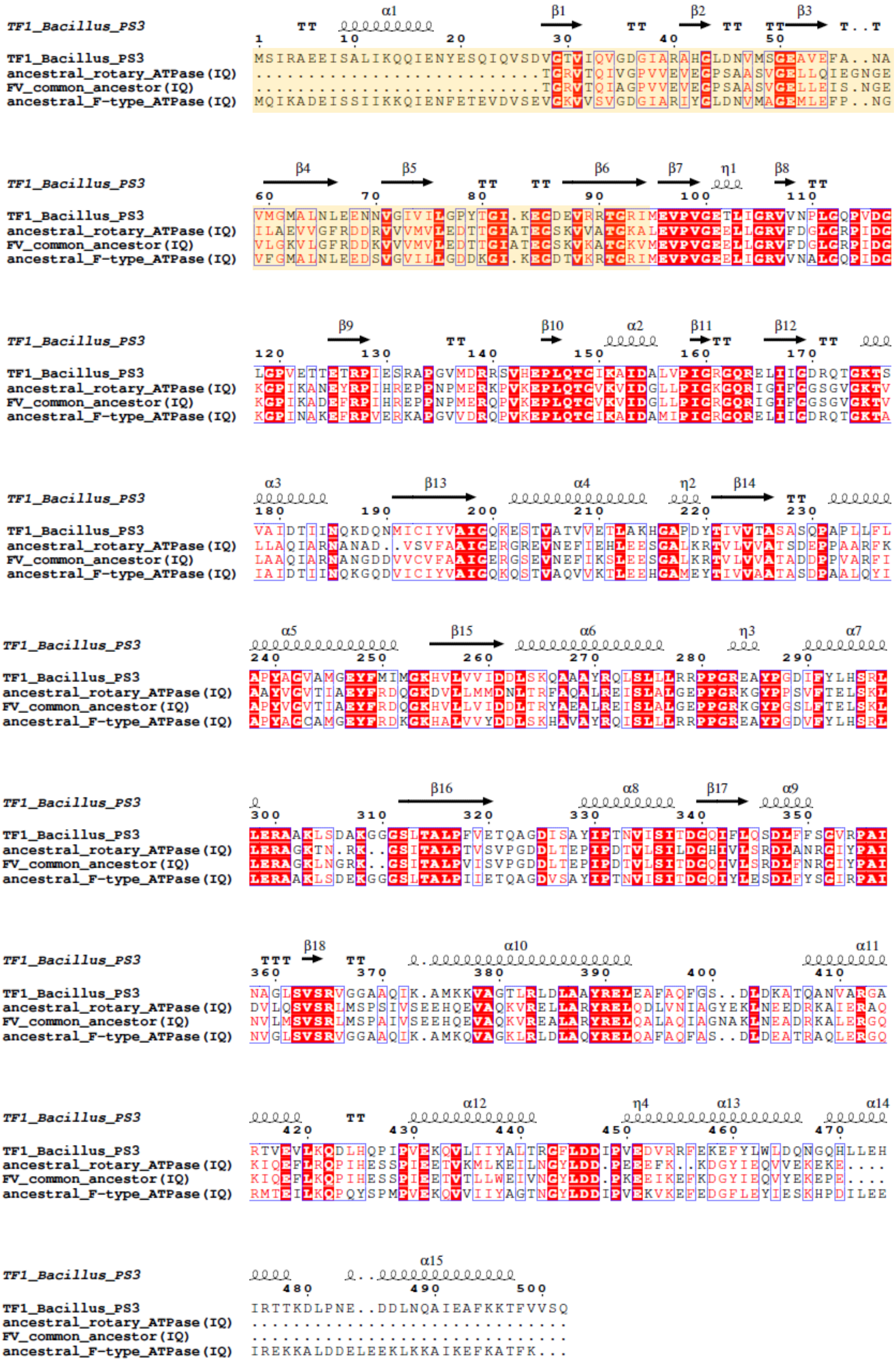
TF1_Bacillus_PS3 TT α1 β1 TT β2 TT TT β3 T..T
1 10 20 30 40 50
TF1_Bacillus_PS3 MSIRAEEISALIKQQIENYESQIQVSDVGTVIQVGDGIARAHGLDNVMSGEAVEFA..NA
ancestral_rotary_ATPase(IQ) ...........................TGRVTQIVGPVVEVEGPSAASVGELLQIEGNGE
FV_common_ancestor(IQ) ...........................TGRVTQIAGPVVEVEGPSAASVGELLEIS.NGE
ancestral_F-type_ATPase(IQ) MQIKADEISSIIKKQIENFETEVDVSEVGKVVSVGDGIARIYGLDNVMAGEMLEFP..NG
TF1_Bacillus_PS3 β4 β5 TT TT β6 β7 η1 β8 TT
60 70 80 90 100 110
TF1_Bacillus_PS3 VMGMALNLEENNVGIVILGPYTGI.KEGDEVRRTGRIMEVPVGETLIGRVVNPLGQPVDG
ancestral_rotary_ATPase(IQ) ILAEVVGFRDDRVVVMVLEDTTGIATEGSKVVATGKALEVPVGEELLGRVFDGLGRPIDG
FV_common_ancestor(IQ) VLGKVLGFRDDKVVVMVLEDTTGIATEGSKVKATGKVMEVPVGEELLGRVFDGLGRPIDG
ancestral_F-type_ATPase(IQ) VFGMALNLEEDSVGVILLGDDKGI.KEGDTVKRTGRIMEVPVGEELIGRVVNALGQPIDG
TF1_Bacillus_PS3 β9 TT β10 α2 β11 TT β12 TT
120 130 140 150 160 170
TF1_Bacillus_PS3 LGPVETTETRPIESRAPGVMDRRSVHEPLQTGIKAIDALVPIGRGQRELIIGDRQTGKTS
ancestral_rotary_ATPase(IQ) KGPIKANEYRPIHREPPNPMERKPVKEPLQTGVKVIDGLLPIGKGQRIGIFGGSGVGKTV
FV_common_ancestor(IQ) KGPIKADEFRPIHREPPNPMERQPVKEPLQTGVKVIDGLLPIGRGQRIGIFGGSGVGKTV
ancestral_F-type_ATPase(IQ) KGPINAKEFRPVERKAPGVVDRQPVKEPLQTGIKAIDAMIPIGRGQRELIIGDRQTGKTA
TF1_Bacillus_PS3 α3 β13 α4 η2 β14 TT
180 190 200 210 220 230
TF1_Bacillus_PS3 VAIDTIINQKDQNMICIYVAIGQKESTVATVVETLAKHGAPDYTIVVTASASQPAPLLFL
ancestral_rotary_ATPase(IQ) LLAQIARNANAD..VSVFAAIGERGREVNEFIEHLEESGALKRTVLVVATSDEPPAARFK
FV_common_ancestor(IQ) LAAQIARNANGDDVVCVFAAIGERGSEVNEFIKSLEESGALKRTVLVVATADDPPVARFI
ancestral_F-type_ATPase(IQ) IAIDTIINQKGQDVICIYVAIGQKQSTVAQVVKTLEEHGAMEYTIVVAATASDPAALQYI
TF1_Bacillus_PS3 α5 β15 α6 η3 α7
240 250 260 270 280 290
TF1_Bacillus_PS3 APYAGVAMGEYFMIMGKHVLVVIDDLSKQAAAYRQLSLLLRRPPGREAYPGDIFYLHSRL
ancestral_rotary_ATPase(IQ) AAYVGVTIAEYFRDQGKDVLLMMDNLTRFAQALREISLALGEPPGRKGYPPSVFTELSKL
FV_common_ancestor(IQ) APYVGVTIAEYFRDQGKHVLLVIDDLTRYAEALREISLALGEPPGRKGYPGSLFTELSKL
ancestral_F-type_ATPase(IQ) APYAGCAMGEYFRDKGKHALVVYDDLSKHAVAYRQISLLLRRPPGREAYPGDVFYLHSRL
TF1_Bacillus_PS3 β16 α8 β17 α9
300 310 320 330 340 350
TF1_Bacillus_PS3 LERAAKLSDAKGGGSLTALPFVETQAGDISAYIPTNVISITDGQIFLQSDLFFSGVRPAI
ancestral_rotary_ATPase(IQ) LERAGKTN.RK..GSITALPTVSVPGDDLTEPIPDTVLSILDGHIVLSRDLANRGIYPAI
FV_common_ancestor(IQ) LERAGKLNGRK..GSITALPVISVPGDDLTEPIPDTVLSITDGQIVLSRDLFNRGIYPAI
ancestral_F-type_ATPase(IQ) LERAAKLSDEKGGGSLTALPIIETQAGDVSAYIPTNVISITDGQIYLESDLFYSGIRPAI
TF1_Bacillus_PS3 TTT β18 TT α10 α11
360 370 380 390 400 410
TF1_Bacillus_PS3 NAGLSVSRVGGAAQIK.AMKKVAGTLRLDLAAYRELEAFAQFGS..DLDKATQANVARGA
ancestral_rotary_ATPase(IQ) DVLQSVSRLMSPSIVSEEHQEVAQKVRELLARYRELQDLVNIAGYEKLNEEDRKAIERAQ
FV_common_ancestor(IQ) NVLMSVSRLMSPAIVSEEHQEVAQKVREALARYRELQALAQIAGNAKLNEADRKALERGQ
ancestral_F-type_ATPase(IQ) NVGLSVSRVGGAAQIK.AMKQVAGKLRLDLAQYRELQAFAQFAS..DLDEATRAQLERGQ
TF1_Bacillus_PS3 TT α12 η4 α13 α14
420 430 440 450 460 470
TF1_Bacillus_PS3 RTVEVLKQDLHQPIPVEKQVLIIYALTRGFLDDIPVEDVRRFEKEFYLWLDQNGQHLLEH
ancestral_rotary_ATPase(IQ) KIQEFLRQPIHESSPIEETVKMLKEILNGYLDD.PEEEFK..KDGYIEQVVEKEKE....
FV_common_ancestor(IQ) KIQEFLKQPIHESSPIEETVTLLWEIVNGYLDD.PKEEIKEFKDGYIEQVYEKEPE....
ancestral_F-type_ATPase(IQ) RMTEILKQPQYSPMPVEKQVVIIYAGTNGYLDDIPVEKVKEFEDGFLEYIESKHPDILEE
TF1_Bacillus_PS3 α15
480 490 500
TF1_Bacillus_PS3 IRTTKDLPNE..DDLNQAIEAFKKTFVVSQ
ancestral_rotary_ATPase(IQ) ..............................
FV_common_ancestor(IQ) ..............................
ancestral_F-type_ATPase(IQ) IREKKALDDELEEKLKKAIKEFKATFK...

**Fig. S5. MSA of Ancestral and Extant F-type ATPase α Subunit Sequences (IQ-TREE)**

The multiple sequence alignment (MSA) includes the sequence and secondary structure of the α subunit from the extant $TF_1$ ATPase, along with ancestral sequences reconstructed based on the IQ-TREE phylogenetic tree. These ancestral sequences comprise the ancestral rotary ATPase ($C^{ancR}$), which represents the common ancestor of the F-type ATPase α/β subunits and the V-type ATPase A/B subunits, the F/V common ancestor ($NC^{F/V}$), which is the common ancestor of the non-catalytic subunits F-type ATPase α and V-type ATPase B, and the ancestral F-type ATPase ($\alpha^{anc}$), which is the common ancestor of the F-type ATPase α subunits. Conserved sequence regions are highlighted in red with white text, while groups of amino acid residues with similar properties, such as hydrophobic/hydrophilic or acidic/basic characteristics, are enclosed in blue boxes. Secondary structure annotations include α-helices (α), β-strands (β), 310 helices (η), and loop structures causing abrupt directional changes in the polypeptide chain (TT), with structural information obtained from the PDB (PDB ID: 6N2Y). The sequence region corresponding to the N-terminal structural foundation involved in complex formation in the hybrid $F_1$ with the ancestral core domains is shaded in orange.

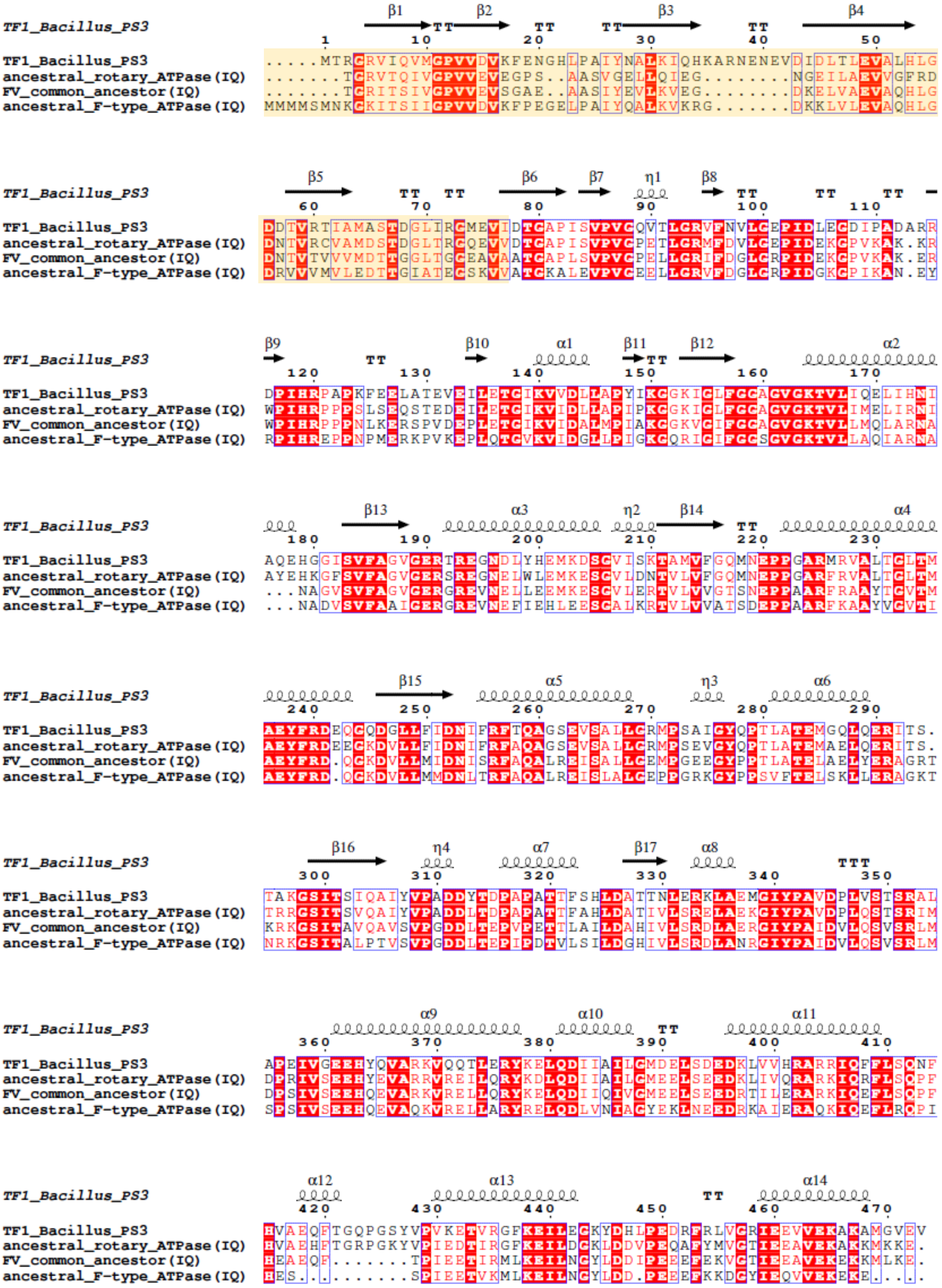

**Fig. S6. MSA of Ancestral and Extant F-type ATPase β Subunit Sequences (IQ-TREE)**

The multiple sequence alignment (MSA) includes the sequence and secondary structure of the β subunit from the extant $TF_1$ ATPase, along with ancestral sequences reconstructed based on the

IQ-TREE phylogenetic tree. These ancestral sequences comprise the ancestral rotary ATPase ($C^{\mathrm{ancR}}$), which represents the common ancestor of the F-type ATPase α/β subunits and the V-type ATPase A/B subunits, the F/V common ancestor ($C^{\mathrm{F/V}}$), which is the common ancestor of the catalytic subunits F-type ATPase β and V-type ATPase A, and the ancestral F-type ATPase ($\beta^{\mathrm{anc}}$), which is the common ancestor of the F-type ATPase β subunits. Conserved sequence regions are highlighted in red with white text, while groups of amino acid residues with similar properties, such as hydrophobic/hydrophilic or acidic/basic characteristics, are enclosed in blue boxes. Secondary structure annotations include α-helices (α), β-strands (β), 310 helices (η), and loop structures causing abrupt directional changes in the polypeptide chain (TT), with structural information obtained from the PDB (PDB ID: 6N2Y). The sequence region corresponding to the N-terminal structural foundation involved in complex formation in the hybrid $F_1$ with the ancestral core domains is shaded in orange.

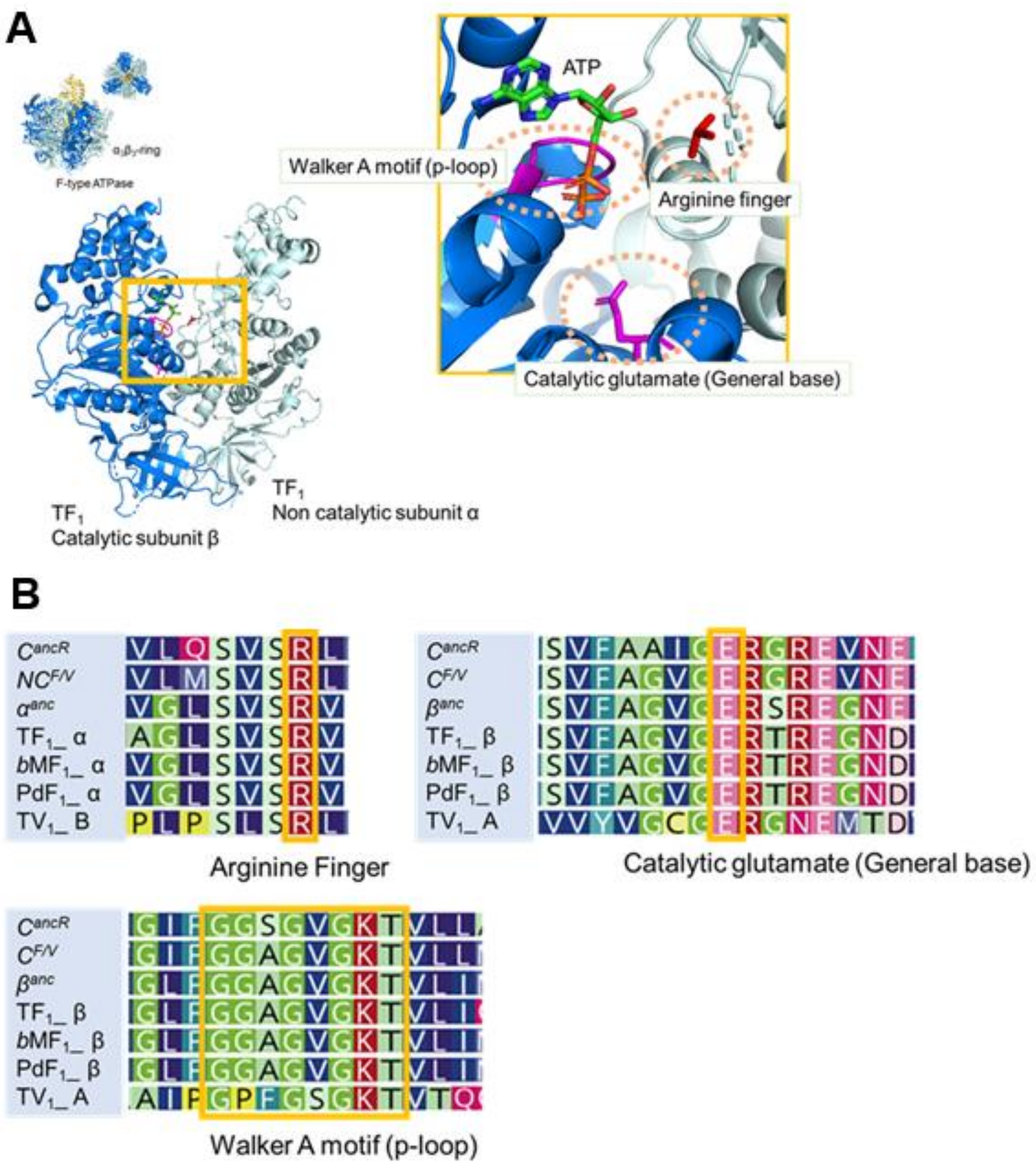


**Fig. S7. Catalytically critical residues and motifs of ancestral sequences**

**A.** Structural location of catalytically critical residues and motifs. **B.** Sequence alignment of the common ancestral sequences and extant sequences. Orange boxes show the catalytic residues and motifs.

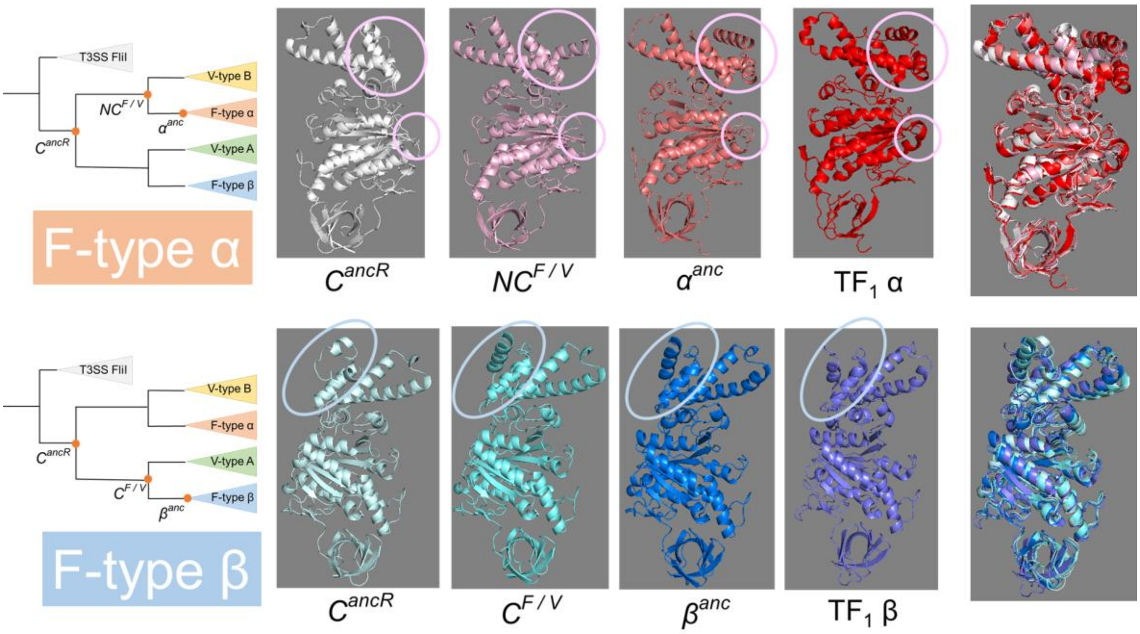


**Fig. S8. Evolutionary Trajectories of F-type ATPase Subunits**

The evolutionary transitions of each subunit from the common ancestor of F-type and V-type ATPases are depicted by arranging ancestral structures sequentially from left to right, corresponding to upstream nodes in the phylogenetic tree. On the right, the structural superpositions of each subunit are displayed. Circular markers indicate regions where significant structural changes were observed. The structure of $TF_1$ is referenced as a representative of extant species. In the figure, the top represents the C-terminal side, while the bottom corresponds to the N-terminal side. Across all ancestral forms and the extant species, the fundamental structural elements remain conserved, including the α-helical structure at the C-terminal region, the β-sheet structure in the central region, and the β-barrel structure at the N-terminal region. The structures of the ancestral types were predicted using AlphaFold2[3], a deep learning-based algorithm developed by DeepMind. The model was implemented using the open-source version of AlphaFold2 with default parameters.

**Table. S2. Proportion of Particles with and without nucleotide**

| | AMP-PNP | | w/o nucleotide | |
|---|---|---|---|---|
| | Number of particles | % | Number of particles | % |
| All particles | 1,753,764 | | 1,357,417 | |
| Hexamer w/ stalk | 97,557 | 5.6 | 398,344 | 29.3% |
| Hexamer w/o stalk | 114,081 | 6.5 | 150,384 | 11% |
| Tetramer w/ stalk | 225,450 | 12.9 | 256,656 | 18.9% |
| Tetramer w/o stalk | 255,786 | 14.6 | 407,761 | 30% |
| Unclassified | 1,060,890 | 60.5 | 144,272 | 10.6% |

### Table. S3. Cryo-EM data collection, refinement and validation statistics

| | #1 Hexamer no stalk (EMDB-49841) | #2 Hexamer with stalk Binding Dwell (EMDB-49839) (PDB 9NVL) | #3 Hexamer with stalk Catalytic Dwell (EMDB-49840) (PDB 9NVM) | #4 Tetramer no stalk Binding Dwell (EMDB-49843) | #5 Tetramer with stalk Binding Dwell (EMDB-49842) |
|---|---|---|---|---|---|
| **Data collection and processing** | | | | | |
| Magnification | 60,000x | 60,000x | 60,000x | 60,000x | 60,000x |
| Voltage (kV) | 300 | 300 | 300 | 300 | 300 |
| Electron exposure (e–/Å$^2$) | 62 | 62 | 62 | 62 | 62 |
| Defocus range (μm) | 0.5-1.5 | 0.5-1.5 | 0.5-1.5 | 0.5-1.5 | 0.5-1.5 |
| Pixel size (Å) | 0.84 | 0.84 | 0.84 | 0.84 | 0.84 |
| Symmetry imposed | C1 | C1 | C1 | C1 | C1 |
| Initial particle images (no.) | 693,389 | 693,389 | 693,389 | 664,417 | 664,417 |
| Final particle images (no.) | 150,384 | 191,889 | 206,455 | 256,656 | 407,761 |
| Map resolution (Å) | 2.77 | 2.47 | 2.54 | 2.70 | 2.46 |
| FSC threshold 0.143 | | | | | |
| **Refinement** | | | | | |
| Initial model used (PDB code) | | 7L1Q | 7L1R | | |
| Model resolution (Å) | | | | | |
| FSC threshold 0.143 | | 2.5 | 3.2 | | |
| Map sharpening *B* factor (Å$^2$) | -67.5 | -57.3 | -62.6 | -80.3 | -72.4 |
| Model composition | | | | | |
| Non-hydrogen atoms | | 24,196 | 24,296 | | |
| Protein residues | | 3,109 | 3,116 | | |
| Ligands | | 6 | 8 | | |
| *B* factors (Å$^2$) | | | | | |
| Protein | | 33.34 | 145.14 | | |
| Ligand | | 44.52 | 147.46 | | |
| R.m.s. deviations | | | | | |
| Bond lengths (Å) | | 0.013 | 0.013 | | |
| Bond angles (°) | | 2.140 | 2.155 | | |
| Validation | | | | | |
| MolProbity score | | 0.91 | 0.92 | | |
| Clashscore | | 0.74 | 1.19 | | |
| Poor rotamers (%) | | 0.78 | 0.54 | | |
| Ramachandran plot | | | | | |
| Favored (%) | | 96.99 | 97.58 | | |
| Allowed (%) | | 3.01 | 2.32 | | |
| Disallowed (%) | | 0.00 | 0.10 | | |

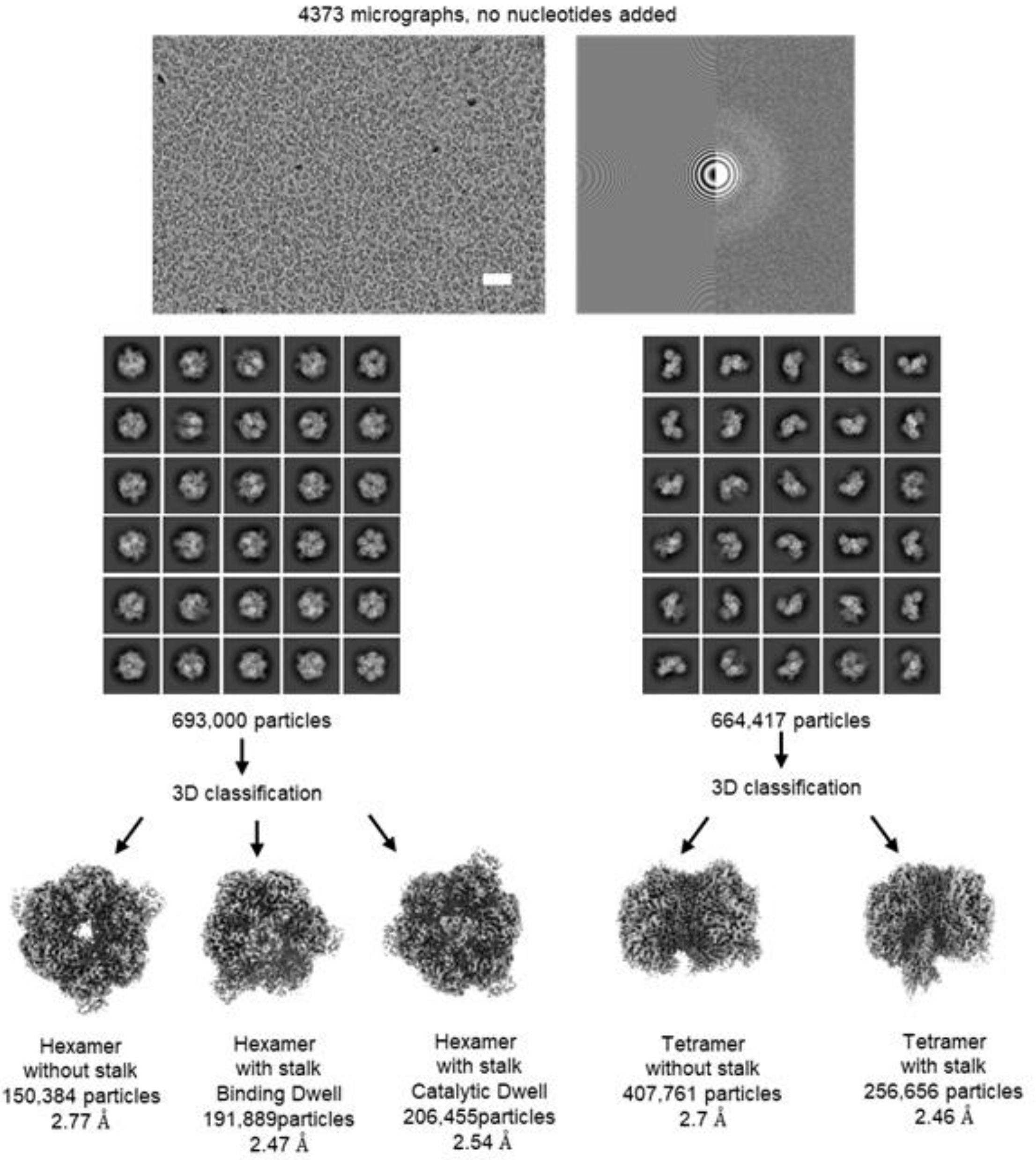


**Fig. S9. Cryo-EM data collection and data processing flowchart**

Micrographs with white scale bar equivalent to 50 nm. 2D and 3D classifications was used to sort the particles into sub-states.

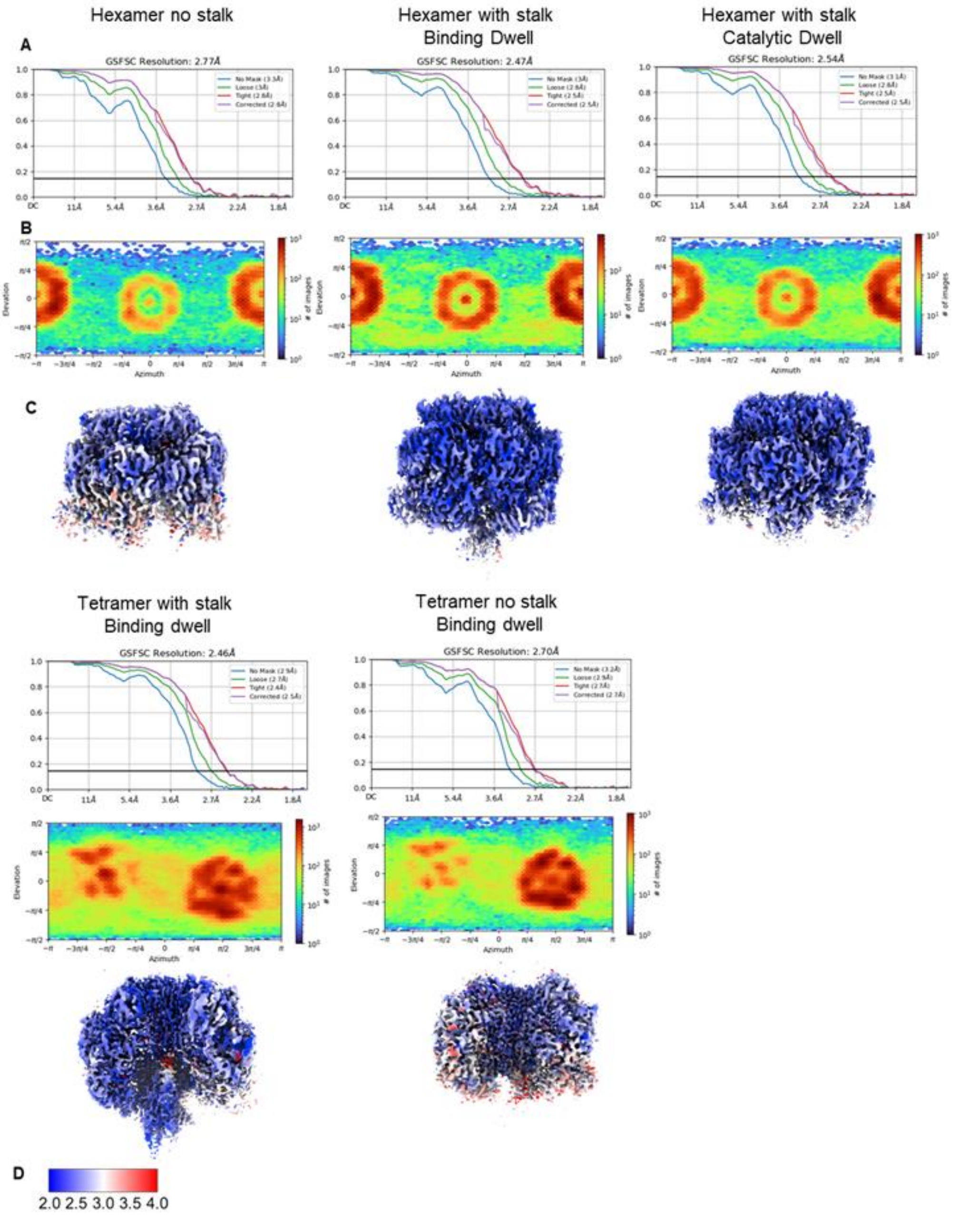


**Fig. S10. 3D FSC Curves and Local Resolution Estimates**

**A.** Gold standard Fourier shell correlation (GSFSC) curves from CryoSPARC. **B.** Viewing direction distribution plot. **C.** Local resolution estimate calculated in cryoSPARC. **D.** Local resolution estimate scale

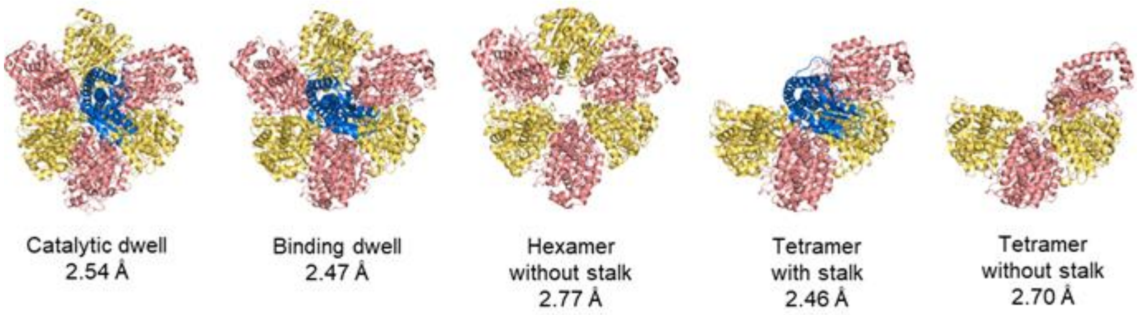


**Fig. S11. Structures of $F_1^{anc\_core}$ obtained in this study**

The structural representation of the common ancestral $F_1$-ATPase is shown with distinct color coding for each subunit: the α-subunit is depicted in pink, the β-subunit in cream, and the γ-subunit in light blue.

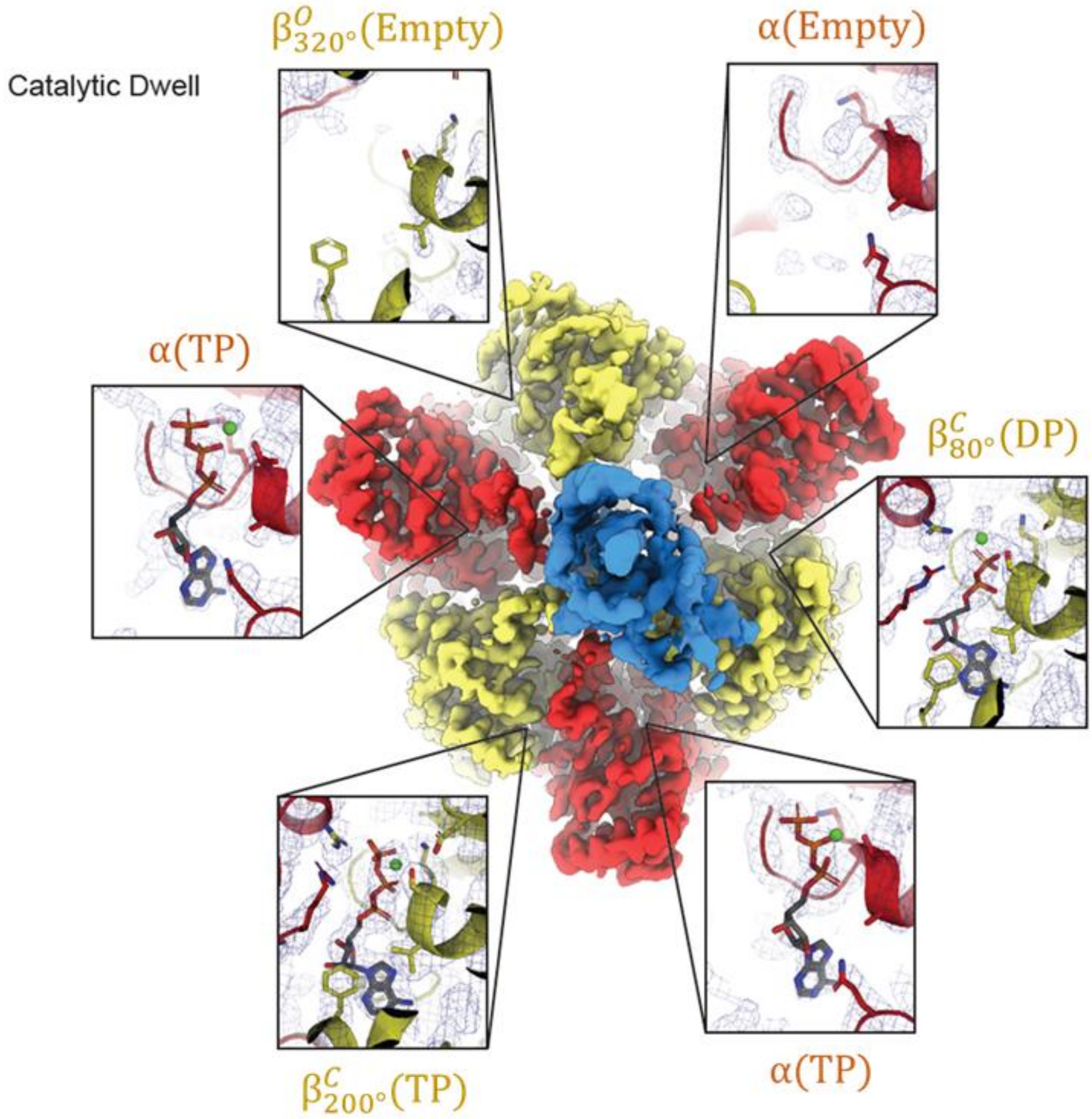


**Fig. S12. Close-up Views of the Nucleotide Binding Sites (Catalytic dwell)**

The cryo-EM maps are shown along with close-up views of each nucleotide binding site. Cryo-EM map is shown as mesh at 4 σ in all close-up panels.

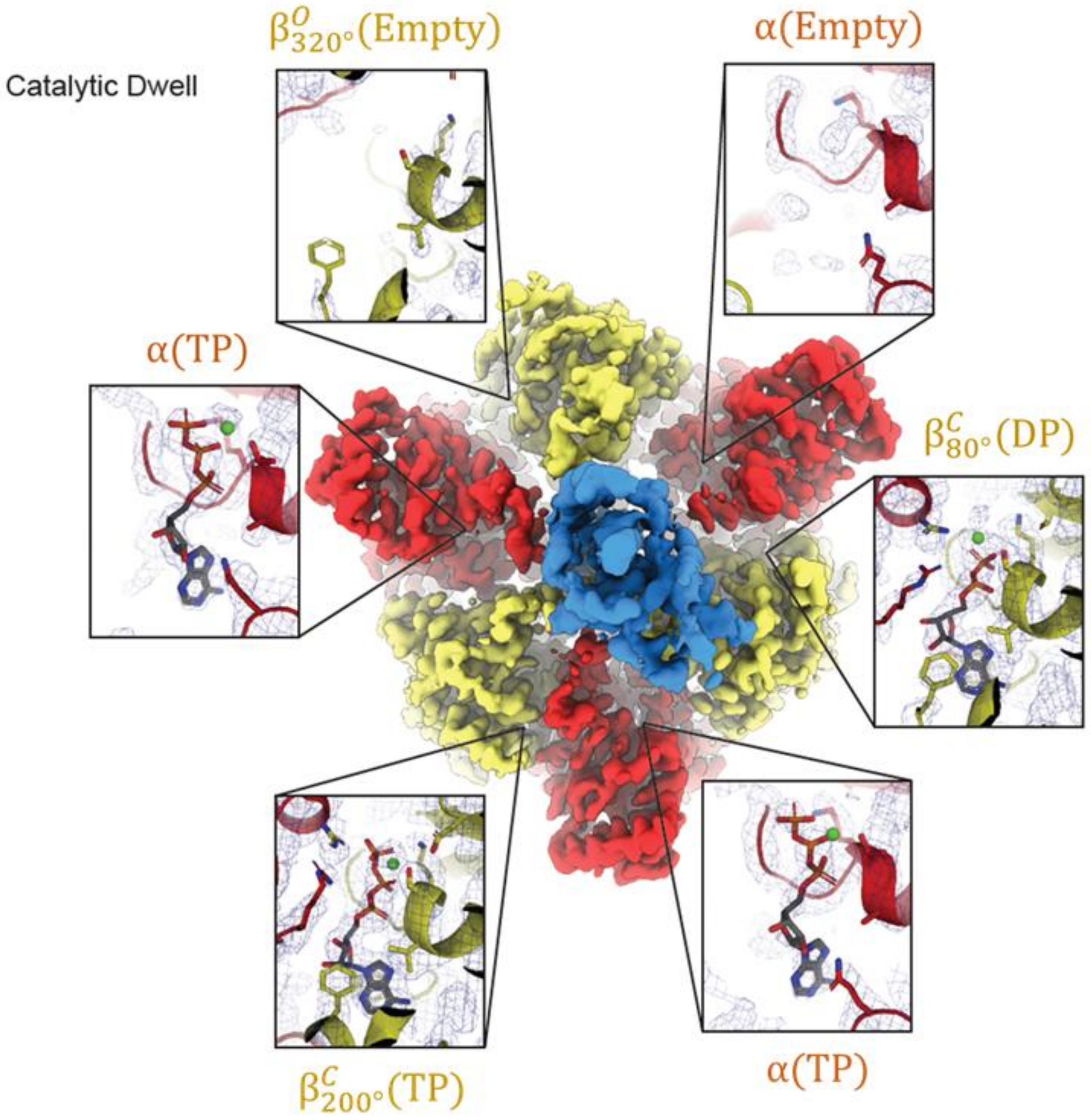


**Fig. S13. Close-up Views of the Nucleotide Binding Sites (Binding dwell)**

The cryo-EM maps are shown along with close-up views of each nucleotide binding site. Cryo-EM map is shown as mesh at 4 σ in all close-up panels.

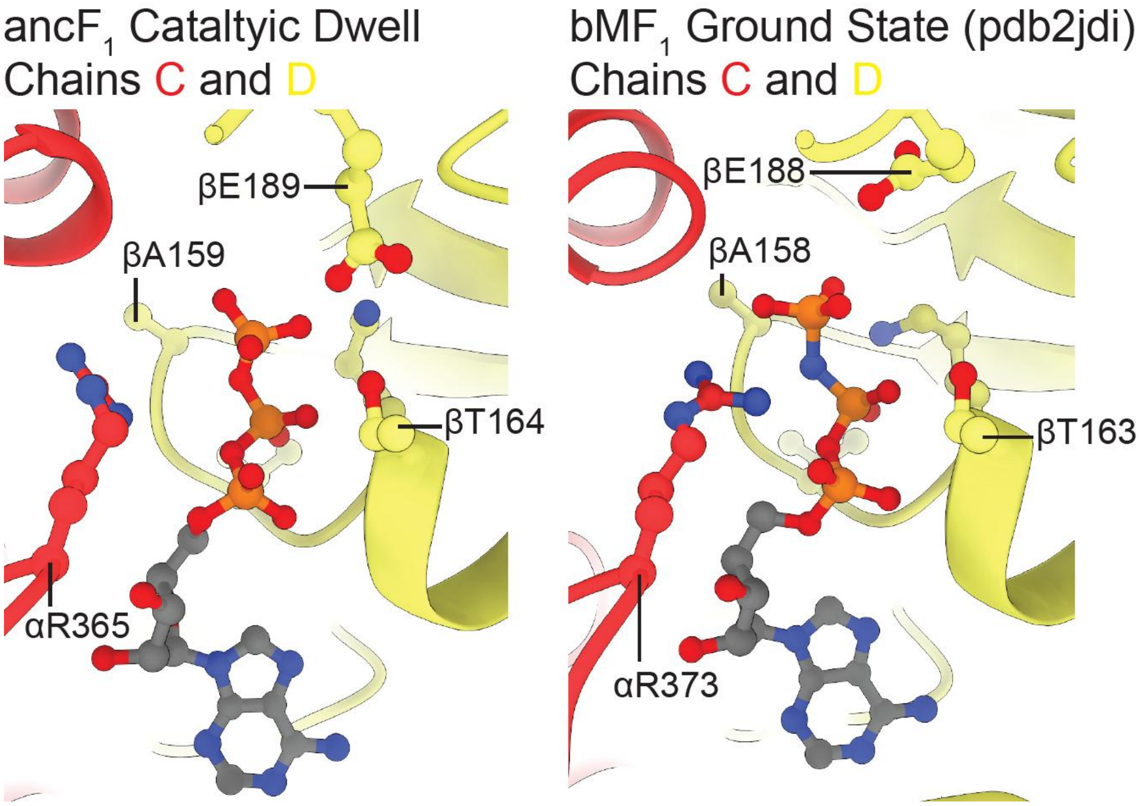


**Fig. S14. Direct comparison between $F_1^{anc\_core}$ and the bovine mitochondrial $F_1$ ground state**

Comparison of the binding site between $\beta^{C}_{200°}$(TP) and α(TP) for $F_1^{anc\_core}$ in catalytic state(left) and bovine mitochondrial $F_1$ in ground state (right). The alpha subunits are colored red, and beta subunits are colored yellow. Residues are labelled to allow for comparison of p-loop (from βA159 to βT164 for $F_1^{anc\_core}$, and from βA159 to βT164 for *b*MF$_1$), arginine finger (αR365 for $F_1^{anc\_core}$, and αR363 for *b*MF$_1$) and catalytic glutamate (βE189 for $F_1^{anc\_core}$, and βE188 for *b*MF$_1$). Although the catalytic glutamate and arginine finger are in slightly different conformations, this is likely just due to the limited resolution of the structural data.

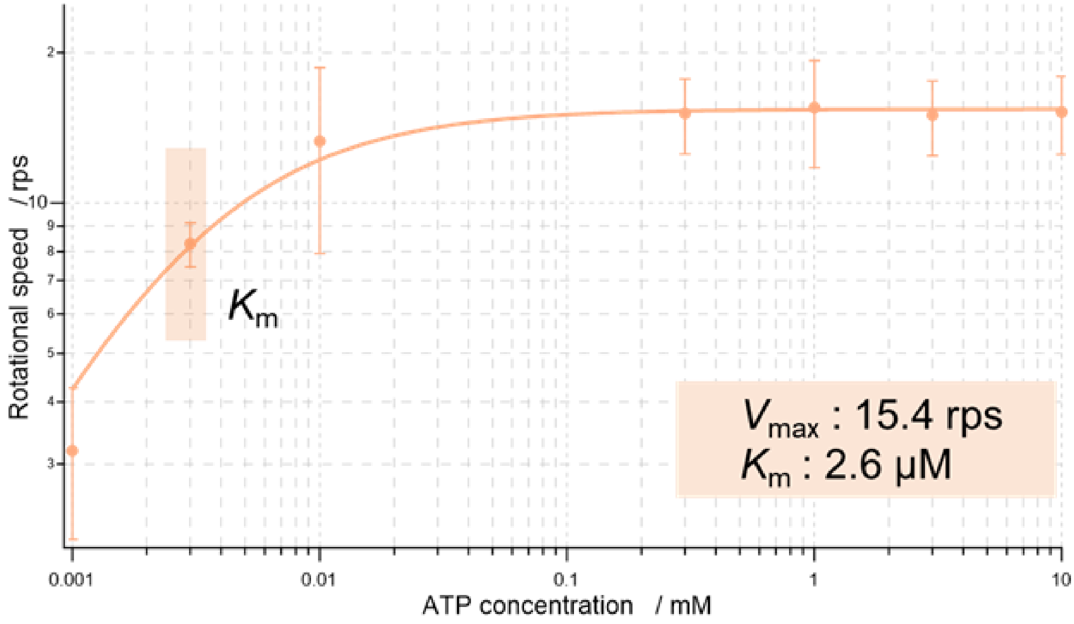


**Fig. S15. Michaelis-Menten Curve of $F_1^{anc\_core}$ for ATP**

The mean rotational velocity of at least three particles was plotted for each ATP concentration and fitted using the Michaelis-Menten equation. Error bars represent the standard deviation (SD). The fitting analysis estimated a maximum velocity ($V_{max}$) of 15.4 rps and a Michaelis constant ($K_m$) of 2.6 μM.

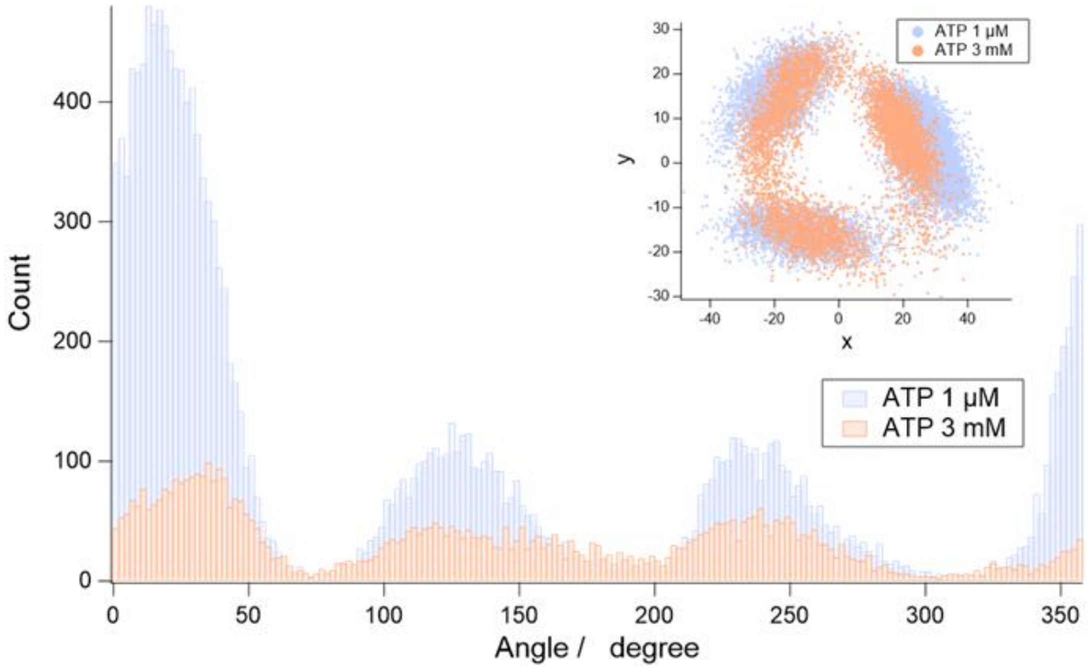


**Fig. S16. Solution Exchange Between 3 mM ATP and 1 μM ATP**

Angle histograms before and after solution exchange. The light purple histogram represents the low ATP concentration condition (1 μM ATP), while the light orange histogram corresponds to the high ATP concentration condition (3 mM ATP). The upper right panel displays x–y plots for each ATP concentration, with color coding consistent with the angle histograms.

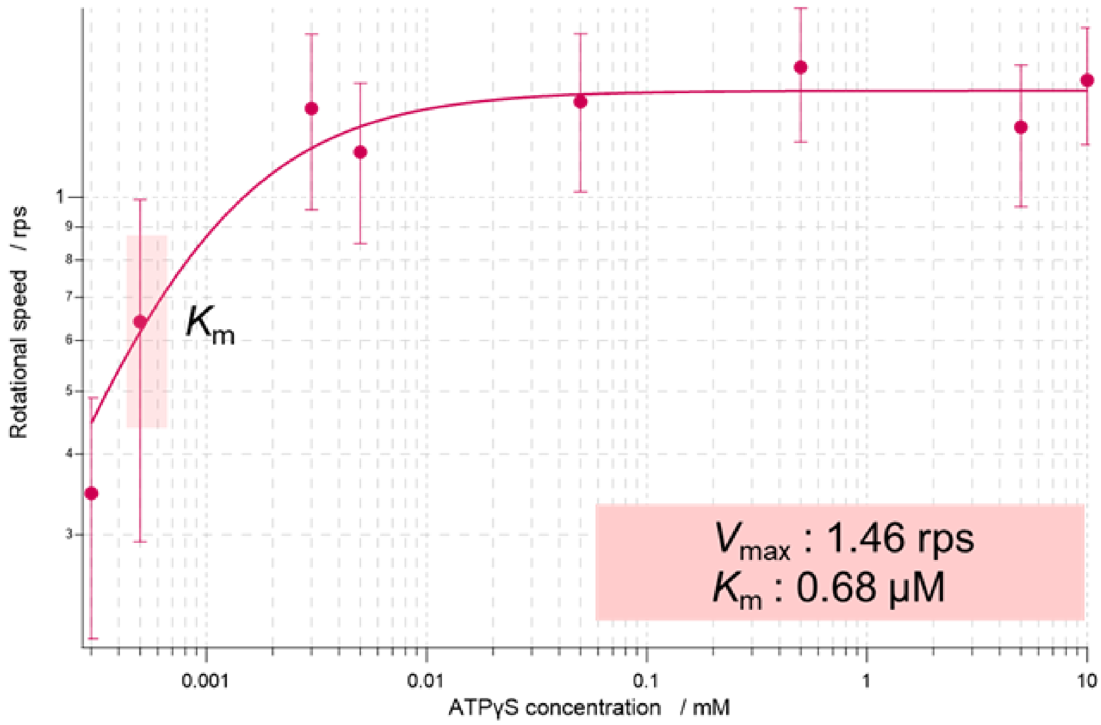


**Fig. S17. Michaelis-Menten Curve of $F_1^{anc\_core}$ for ATPγS**

The mean rotational velocity of at least three particles was plotted for each ATPγS concentration and fitted using the Michaelis-Menten equation. Error bars represent the standard deviation (SD). The fitting analysis estimated a maximum velocity ($V_{max}$) of 1.46 rps and a Michaelis constant ($K_m$) of 0.68 μM.

**Table. S4. $Q_{10}$ Factor of $F_1^{anc\_core}$**

| Temperature | w/ LDAO | w/o LDAO |
|---|---|---|
| 20°C - 30°C | 1.33 | 2.91 |
| 25°C - 35°C | 1.60 | 1.89 |
| 30°C - 40°C | 1.63 | 1.79 |
| Average | 1.52 | 2.20 |

The ATPase activity of $F_1^{anc\_core}$ was measured at various solution temperatures to estimate the $Q_{10}$ factor, which represents the fold increase in ATPase activity upon a 10°C rise in temperature. ATPase activity measurements were conducted twice at each temperature, and the average values were used to estimate the $Q_{10}$ factor.

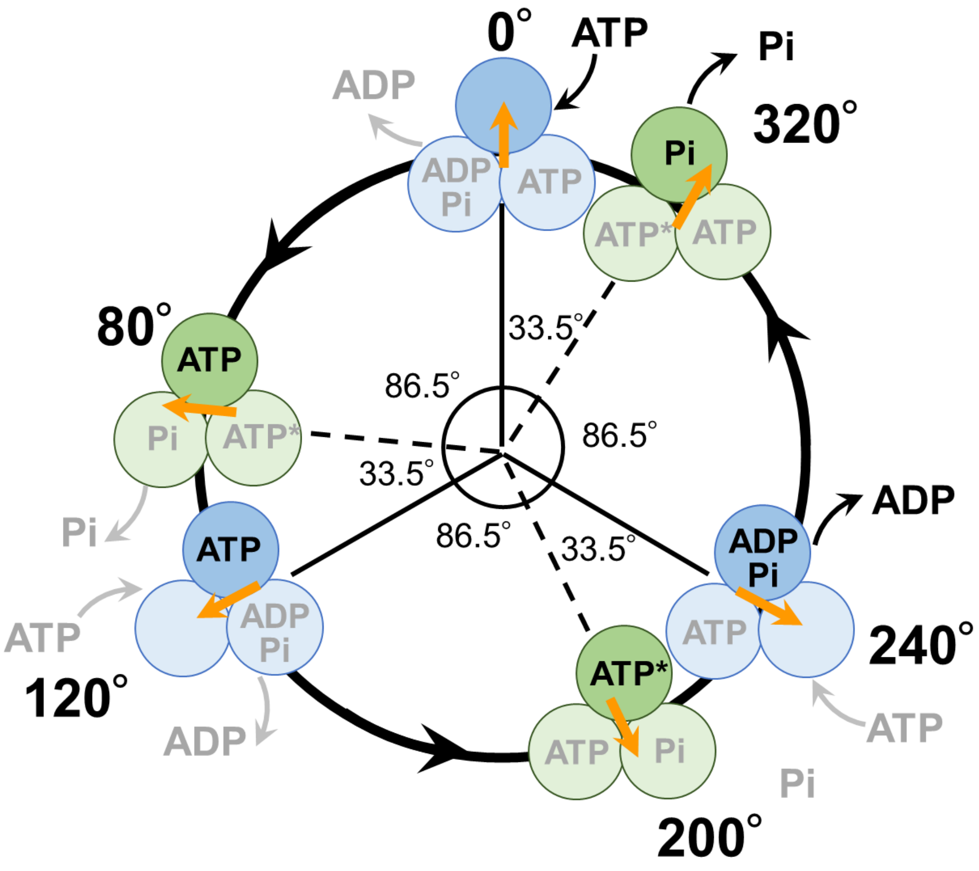


**Fig. S18. Rotary Catalytic Mechanism of Hybrid $F_1$ with the Ancestral Core Domains**

The three circles represent the three β subunits of the hexameric ring, with the nucleotides bound at each β subunit's binding site indicated inside the circles. ATP* denotes the activated state of ATP prior to hydrolysis. The orange arrow indicates the rotation of the γ subunit. The blue and light blue markers represent the binding dwell positions, while the green and light green markers indicate the catalytic dwell positions, where rotation pauses.

Due to the significantly faster reaction kinetics occurring at the catalytic dwell compared to the binding dwell, the rotational angles of the binding dwell are represented by solid lines, whereas those of the catalytic dwell are depicted by dashed lines. The difference in rotational angles between the binding dwell and catalytic dwell corresponds to the angular shift revealed by cryo-EM structural analysis.
